# Neural Correlates of Perceptual Decision Making in Primary Somatosensory Cortex

**DOI:** 10.1101/2025.03.29.646003

**Authors:** Alex G. Armstrong, Yurii Vlasov

## Abstract

The brain is thought to produce decisions by gradual accumulation of sensory evidence through a hierarchically organized feedforward cascade of neuronal activities that transforms early stimulus representations in the primary somatosensory cortex (S1) to a perceptual decision processed in pre-motor areas. Recently, this prevailing view has been challenged by observation of choice- correlated neural activity as early in the hierarchy as S1. Here, to reconcile these seemingly controversial observations, we employ ethological whisker-guided navigation of mice in a tactile virtual reality paradigm combined with dense electrophysiological recordings in whisker-related wS1. Leaving only a pair of C2 whiskers for mice to navigate with, we effectively designed an information bottleneck for sensory input to decision making. We show that neural activity during sensory evidence accumulation exhibits dramatic collapse of the high-dimensional spiking activity to just a single latent variable followed by a slower and almost synchronous ramping up across the whole cortical column. We show that this variable is consistent with models of gradual accumulation of noisy sensory evidence to a decision bound. These observations indicate that S1 may directly participate in a categorical coding of all-or-none decision variable via cortico-cortical feedback loops through which sensory information reverberates to be transformed into perception and action.

**Significance Statement:** By employing ethological whisker-guided navigation of mice in a tactile virtual reality paradigm combined with dense electrophysiological recordings in whisker-related wS1, we show that neural activity during sensory evidence accumulation exhibits dramatic collapse of the high-dimensional spiking activity to just a single latent variable followed by a slower and almost synchronous ramping up across the whole cortical column. We show that this variable is consistent with models of gradual accumulation of noisy sensory evidence to a decision bound. These observations indicate that wS1 may directly participate in a categorical coding of all-or-none decision variable. Naturalistic perceptual decisions during active whisker-guided navigation High-dimensional activity in S1 collapses to a single variable prior to decision This ramping spiking activity is consistent with drift-diffusion decision models Indicates direct involvement of S1 in evidence accumulation

## INTRODUCTION

The prevailing view on perceptual decision making mostly adapted from influential research in the visual system of primates has been that sensory flow is gradually transformed from stimulus representations in the primary somatosensory cortex (S1)^1^ to neural activity correlated with decision in the secondary somatosensory cortex^2^ (S2) up to a categorical all-or-none perceptual decision computed in pre-motor areas^3,4^. Here, close to the top of the hierarchy, internal decision variables are accumulated in a form of ramping neural activity^3–6^ that is transformed into a decision when a decision boundary is reached^7^. Serial transformation of sensory inputs along bottom-up hierarchical pathways has been observed in rodents engaged in whisker-discrimination tasks^8,9^ with an increasingly stronger association with perceptual choice in S2^10,11^ and pre-motor areas^12–14^ rather than whisker-related wS1^9^. The function of wS1 is usually analyzed^15^ in the context of extracting complex, high-order features of sensory input^16–18^. It has been observed, however, that wS1 may integrate sensory information over time^19^ and contribute to goal-directed behavior^20,21^. More recent experiments^22–25^ have shown neural activity in subpopulations of layer 2/3 of wS1 neurons is correlated with behavioral choice. In particular, wS1 neurons projecting to S2 were found to increase their spike rate, predicting the choice^22^. Correlation with choice was found to be strongest for the feedback S2 to wS1 projecting neurons^23^. Hence these new findings, although seemingly contradicting earlier views, emphasize the role of feedback loops through which sensory information reverberates^26^ before transforming into perception and action.

Most decision-making studies in rodents, and in part, primates, rely on stereotyped behavioral tasks requiring intensive weeks-long training to obey artificial decision rules that may affect underlying neural computations. To get further insights into processes of transformation of sensory inputs into perception and decision, it is crucial^27^ to employ behavioral tasks that are closer to animals’ behavior in their natural habitat, which brain circuits were specialized to complete during evolution. Animal decision making is often associated with fast dynamics of navigating terrain, outrunning predators, or pursuing prey that echoes fast ms-scale dynamics of “reverberations” of sensory information along feedback loops. While 2-photon microscopy used in previous studies^10,11,17,22–25^ can provide access to thousands of neurons simultaneously, it is typically limited to shallow cortical layers and difficulty resolving fast neural dynamics.

Here, addressing these challenges, we harnessed the natural whisker-guided navigation of mice using tactile virtual reality^17,28,29^ combined with dense multielectrode array electrophysiological recordings. Head-fixed mice navigate a virtual corridor at elevated speeds, defined by a pair of motorized walls, using only their whiskers, closely resembling navigation in their natural habitat^28^. As opposed to traditional designs, our task is untrained and unrewarded. Leaving only a pair of C2 whiskers for mice to navigate with, we designed an information bottleneck for sensory input to decision making^29^, strategically placing a microelectrode probe into the C2 principal whisker barrel and aligning it perpendicular to the cortical layers, recording neural activity across entire cortical depth. This design reveals unexpectedly strong correlation of neural activity across the columnar circuits with decision variables. In fact, during decision formation, variance of the high-dimensional spiking activity collapses to just a single latent variable, while spiking synchronously ramps up across the cortical column. We show that this variable is consistent with models of gradual accumulation of noisy sensory evidence to a decision bound^5,7,26,30^. These observations indicate that primary sensory cortex not only retains relatively weak and sparce choice-related activity, as it was shown previously^10,11,19–25^, but also, in our naturalistic behavioral task, directly participates in a categorical coding of all-or-none decision variable into which the majority of neural populations are recruited.

## Results

### Ethological decision-making during single-whisker-guided navigation in virtual reality

Loading cortical circuits with difficult yet manageable cognitive tasks and at the same time elicit ethologically relevant behavior, we designed a naturalistic tactile virtual reality^17,28,29^ paradigm (SI Methods) where head-fixed mice run on a 3D treadmill, while navigating using only their whiskers (**Fig.1A**). Animal’s movement in lateral (V_lat_) and forward (V_for_) directions is used to provide positive feedback to movement of the walls (V_wall_) positioned at both sides of the snout, generating a closed-loop (CL) virtual reality (VR) corridor for the animal to follow (**Fig.1B**). Importantly, all but the C2 whiskers (second longest vibrissae oriented parallel to the ground) are trimmed, restricting sensory information flow into the primary somatosensory cortex.

**Figure 1.**
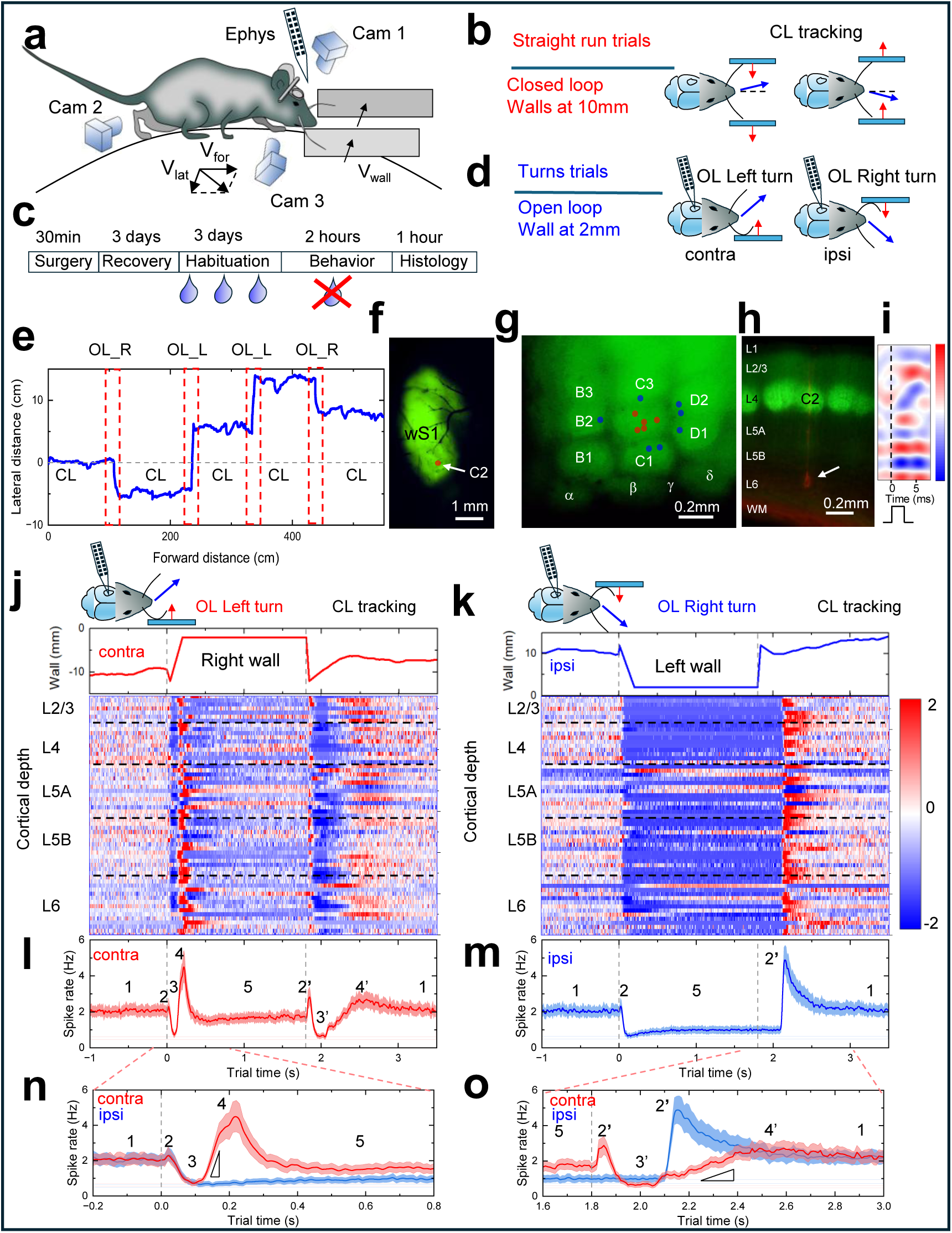
Neural activity in principal whisker barrel column exhibits strong, staged, and synchronous modulation during decision-making. **A)** Schematics of VR navigation experiments. Forward Vfor and lateral Vlat speeds of animal on the suspended sphere are used to calculate positive feedback signal Vwall that is fed into the motorized walls to approach or retract from the animal’s snout. Additional cameras are used to track animals’ legs as well as whisker interactions with the wall. **B)** Closed loop (CL) experiment when animal is navigating a straight VR corridor the distance between the wall and snout is 10mm with walls following the animal’s trajectory in positive feedback closed loop. **C)** Protocol includes 2-3 days habituation in CL VR navigation as shown in B) when animal is rewarded for running at elevated speeds. Followed by a single 2-hour terminal experiment where unrewarded 2AFC task with walls approaching the snout at 2mm distance are introduced in an open loop. **D)** Open loop (OL) 2AFC task with right (contra-lateral to ephys probe) or left (ipsilateral) walls quickly approach the snout forcing animal to turn left (OL_L) or right (OL_R). **E)** Example trajectory of animal navigating a randomized sequence of OL_L and OL_R turns separated by CL straight sections. **F)** Fluorescence micrograph after headbar attachment surgery with visible barrel cortex used for precise in vivo targeting of a principal whisker barrel. **G)** Barrel cortex in fixed brain post hoc. Locations of the probe in the principal (red) and neighboring (blue) whisker barrels reconstructed from histology. **H)** Superimposed fluorescence micrographs of a coronal slice through principal whisker barrel column showing a probe track (red) and a microlesion (arrow) used for reconstruction of electrode locations. **I)** Current-source density (CSD) trace following optogenetic activation used to register the location of layer 4 along the probe. **J)** Top: Right wall position during the OL_L turn. Bottom: Trial-averaged z-scored spiking activity of 65 individual simultaneously recorded units across the principal barrel column for 266 OL_L left turns. Units ordered by cortical depth and assigned to layers. **K)** Top: Left wall position during the OL_R turn. Bottom: Same as J) for 265 OL_R right turns. **L)** Population-averaged spike rate for data in J) for OL_L left turns. Numbers correspond to modulation epochs 1 through 5. Red curve is population mean. Red error band corresponds to s.e.m. Vertical line corresponds to start of wall movement. **M)** Same as L) for data in K) for OL_R right turns. **N)** Population-averaged spike rates from L) and M) for first 1s of turn. Vertical line corresponds to start of wall movement. Red/blue curves correspond to left/right turns. Triangle shows linear fitting of the slope. **O)** Same as N) but for last 1.4s of turn trials. Triangle shows linear fitting of the slope.

Typically, animals require only 3 days of habituation sessions with CL straight trials (**Fig.1C**) to acclimatize to head-fixed locomotion (SI Methods) with water rewards given only to encourage running at fast speeds (21±4cm/s, mean ± s.d., **Fig.S1A**). When habituated, animals undergo a single terminal 2-hour long behavioral session that is unrewarded (**Fig.1C**). To generate an analogous two-alternative forced choice (2AFC) paradigm, we introduce an oddball deviant stimulus (open loop (OL) turn trials, **Fig.1D**). After a baseline CL straight section, the feedback signal is terminated, and walls are briefly (within 10ms) removed beyond the reach of the C2 whiskers to produce consistent sensory input across trials. Then just one wall approaches the snout (within 0.2s) and stays at a pre-defined constant distance for 1.8 seconds before retracting again (**Fig.1D**). Sensing fast-approaching obstacles with their C2 whiskers, animals are forced to change running direction (OL left or right turns, **Fig.1D**) to avoid the expected collision. Typical 20s long trajectory of the animal locomotion is presented in **Fig.1E** exhibiting sections of straight runs (CL) and randomized left (OL_L) and right (OL_R) turns performed while animal is running at elevated speed.

We previously showed^29^ that local ablation of layer 4 (L4) of the principal C2 whisker barrel column, but not neighboring barrels, results in the inability of mice to produce turns guided by whisker contra-lateral to the lesion, while still performing turns guided by ipsi-lateral whisker. This is consistent with previous reports on impairment of whisker-guided gap crossing after sub-columnar micro-infarction^31^. Thus, we created a bottleneck of sensory information critical for making decisions in our untrained and unrewarded 2AFC task.

### Population dynamics in C2 barrel column exhibit strong, staged, and synchronous modulation during decision-making

For electrophysiological studies (SI Methods, **Fig.S2**), either 64- or 960- channel (Neuropixel 1.0) multielectrode array probe (Supplementary information **(SI) Table 1**) was inserted into C2 principal barrel column (SI Methods, **Fig.1F, G, H**) aligned perpendicular to cortical layers. *Scnn1a-TG3-Cre* x *Ai32* transgenic mice enabled targeting of principal barrel without stereotaxic surgery and enabled optogenetic tagging (SI Methods). *In-vivo* fluorescent images during surgery (**Fig.1F**) guide the insertion (**Fig.S3A, B**). *Ex-vivo* histological mapping of barrel cortex (**Fig.1G, Fig.S3C, D**) and coronal slices (**Fig.1H, Fig.S3E-H**) together with *in-vivo* optogenetic tagging and corresponding current source density (CSD) map (**Fig.1I**) confirmed probe placement and electrode assignment (SI Methods).

Trial-averaged population activity (SI Methods) for example animal (#489, **SI Table 1**) is shown in **Fig.1J** and **Fig.1K** (other examples in **Fig.S4**) for OL_L left turns, where right wall approaches the snout while recording in the left hemisphere (contralateral turns, **Fig.1D**), and for OL_R right turns where left wall approaches the snout instead (ipsilateral turns, **Fig.1D**), correspondingly. Spiking activity across all cortical layers is strongly modulated during approach (t=0s) and retraction (t=1.8s) of the walls and can roughly be divided into 5 consecutive epochs as shown in population-averaged (SI Methods) neural activity plots (**Fig.1L, M**). At the start of the turn when the wall is fast approaching (**Fig.1N**), a stable spiking (t<0s, 2±0.1Hz, mean ± s.d.) of epoch 1 is disrupted by a sharp peak of epoch 2 (t=0.026s), much stronger pronounced in single L4 unit activity (**Fig.S5A, B**) for both ipsi- and contra-lateral turns. Epoch 3 follows, exhibiting strong suppression of spiking activity down to 0.7±0.1Hz (mean ± s.d.) for both left and right turns (**Fig.1N**). While for ipsilateral turns the suppression lasts until the turn end (t=1.8s, blue curve **Fig.1O**), contralateral turns produce strong ramping activity, epoch 4,(t=0.25s, red curve in **Fig.1N**) that is quenched at longer times to a stable epoch 5 until the turn end (t=1.8s, red curve in **Fig.1O**). Here, similar staged modulation is observed (**Fig.1O**) when both contra-lateral and ipsi-lateral walls move back to 10mm distance to resume CL wall tracking (at t>2s). Importantly, all 5 epochs are major features that dominate the activity of each 65 simultaneously recorded units (single unit examples in **Fig.S5**), highly reproducible across 531 randomized trials with jitter of just 30ms.

Overall, both single unit and population-averaged spike rates represent strong, staged, and almost synchronous modulation across the principal barrel column. Such synchronization of firing across the cortical column is commonly observed ^32^ during brief whisker stimulation^33–35^, as well as in decision making that is attributed to top-down feedback regulation^36,37^ influencing experience-derived predictions^10^, establishment of attentional state^38^ and working memory^39^. In what follows, we argue these distinct epochs of neural modulation are strongly correlated with different behavioral stages of the decision-making process.

### Behavioral correlates of decision-making mark duration of a choice period

To extract behavioral measures and correlates of decision making from start of choice-relevant sensory input, motor action execution of the choice, associated reaction times (RT), and psychometric curves, we analyzed the statistics of animal trajectories, velocities, and accelerations (n=7). Trajectories for 414 consecutive randomized left and right turn trials for an example animal (#503, other examples in **Fig.S6A, B**) are shown in **Fig.2A**. While straight CL section produces trajectories tightly confined within 0.14cm±0.55 (mean ± s.d.) (**Fig.2B**, bottom histogram at t=0s), distributions of lateral distances after the turn widen over time (**Fig.2B**, middle t=1s, and top t=3s) indicating gradual accumulation of navigation uncertainties. Defining threshold for correct trials as ±1cm lateral clearance at t=1s, the success rate reaches 95% (14 failed red trajectories in **Fig.2A**). To measure psychometric curves, additional experiments were performed (n=7, SI Methods, **Fig.S6E- G**) with lateral distance in the OL turn trials varied from 8mm (whisker barely touching the wall) to 1mm (whisker strongly bent by the wall). The resulting success rate as a function of wall distance (**Fig.2C**) exhibits a strong increase towards shorter distances. Better threshold clearance is achieved by steeper turn angles (**Fig.2D**) at closer wall distances^28^ reflecting the stronger evidence for the animal to turn faster to avoid anticipated collisions^29^.

**Figure 2.**
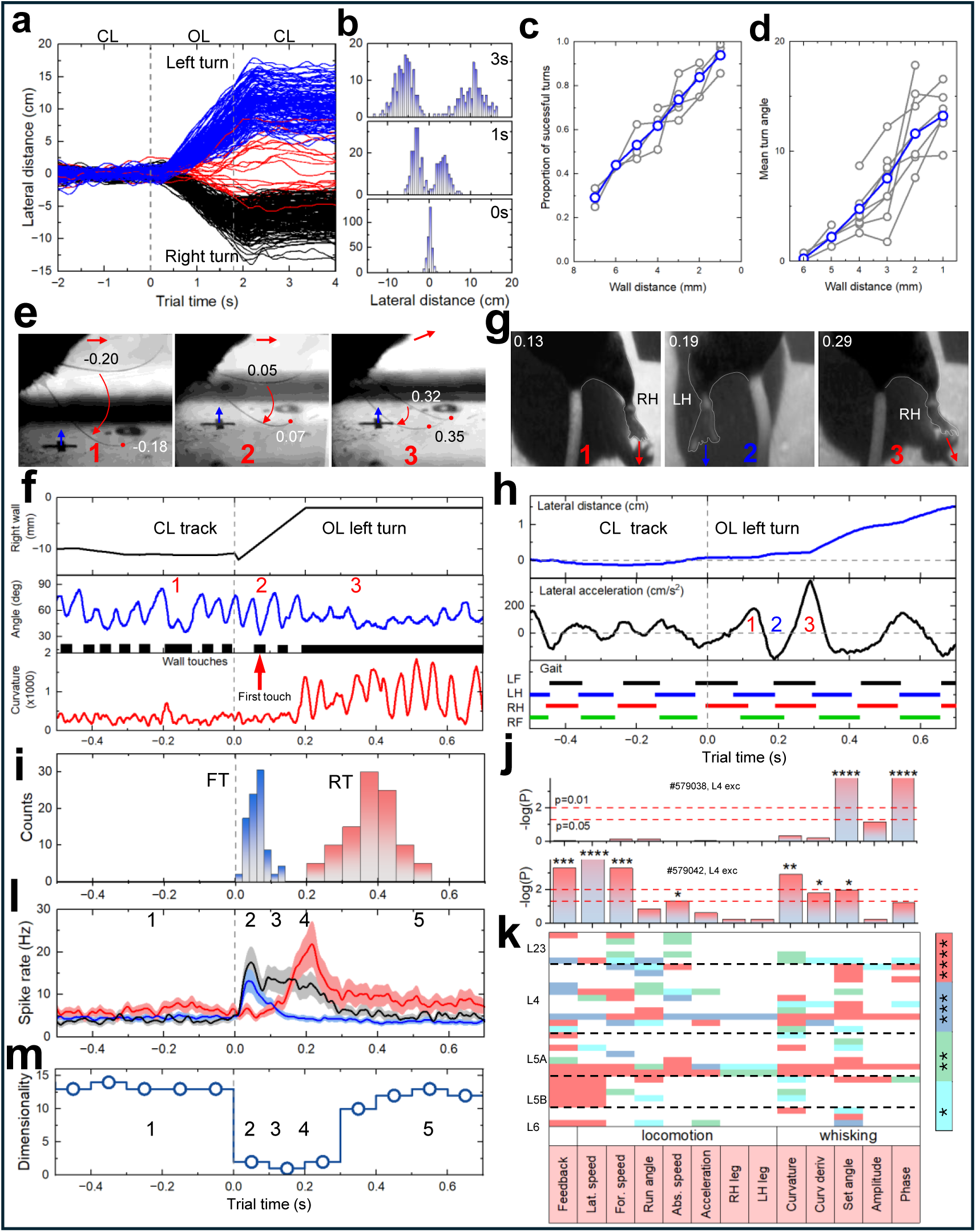
Stages of decision making strongly correlate with stages of neural activity modulation. **A)** Trajectories for 414 trials randomized between OL_L (blue) and OL_R (green) turns. Red curves correspond to error trials. Vertical lines correspond to histograms in B). **B)** Lateral distance distributions at t=0s (bottom), t=1s (middle), and t=3s (top). While at the beginning of a turn the lateral distribution is narrow, they broaden, indicating gradual accumulation of navigation uncertainties. **C)** Percentage of success trials as a function of the wall distance during OL experiments. Psychometric curves measured for left and right turns (n=4, gray, 20+ trials per distance) and population average (blue). **D)** Mean turn angle for the same data as C) (n=4). **E)** Frames from high-speed video of Cam1 (Video S1) showing whisking during CL straight run (left), first touch to approaching wall (middle), and during OL interaction during turn (right). Black bars correspond to whisker touching wall. Red arrow shows direction of mouse run; blue arrows show wall movement. Numbers correspond to the same in F) **F)** Top: Position of the right wall during left turn. Middle: whisker angle. Black bars identify whisker touching the wall, with first touch to approaching wall marked by arrow. Bottom: Same for whisker curvature. Vertical line corresponds to start of wall movement. **G)** Frames from video taken by Cam2 (Video S1) tracking positions of hind legs during 3 consecutive strides corresponding to decision execution. Red arrows show direction of push. Timestamps and numbers correspond to angle change maxima and strides in H). **H)** Top: Lateral distance during left turn trial. Middle: Angle change curve with stereotypic RH leg push at beginning of turn corresponding to G) at t=0.37s. Bottom: tracked legs following diagonal trotting gait pattern. **I)** Histograms of first whisker touch (FT, blue) and reaction time defined by stereotypic RH leg push (RT, red) extracted from recordings similar to F) and H). **J)** Example units used in multiparameter generalized linear-nonlinear Poisson model (GLM) fitting of spiking activity during 55 CL straight run trials. P values are from likelihood ratio test between full and reduced model (SI Methods) to determine which variables are significantly encoded by each unit. **K)** Matrix showing GLM results for 30 units, organized by cortical layers. Color is level of significance from log likelihood ratio test, revealing which variables are encoded by which units. **L)** Population averaged spike rate for OL_L turn trials when animal was sitting for periods of 2s so that active whisking was maintained (20 trials, black) and running (96 trials, red). Blue curve is for OL_R trials for reference. Error band corresponds to s. e. m. Vertical line corresponds to start of wall movement. Numbers are neural epochs. **M)** Intrinsic dimensionality of the neuron population during OL_L turn trial. Dimensionality refers to number of latent variables required to explain at >90% of variance of neural data, showing collapse to single latent variable during choice. Numbers are neural epochs.

The contra-lateral (right) C2 whisker (**Fig.2E**) is tracked with high-speed overhead camera (Cam1 in **Fig.1A**, 1000fps) while interacting with the approaching wall snout. Whisker angles (**Fig.2F**, middle) curvatures (**Fig.2F**, bottom), and touch times (**Fig.2F**, black bars) are extracted from videos (SI Methods, **Fig.S11**). Whisking during CL straight run (t<0) is executed at 15±2Hz (mean ± s.d.) with an amplitude of just 39°±5 (mean ± s.d.) around a set point of 58±5° (mean ± s.d.), typical for “foveal” whisking during fast locomotion^8,28^. During foveal whisking, a whisker periodically palpates the surface during short touches (**Fig.2F**, black bars) occurring mostly during retraction cycles. When the wall is close, it restricts whisking amplitude (whisker angle clamped at 68°±12 (mean ± s.d.) for t>0.2s in **Fig.2F**, middle). However, curvature is still strongly modulated (**Fig.2F**, bottom) indicating whisking remains intact. The first touch (FT) of the whisker to approaching wall is detected at t=0.07s (red arrow **Fig.2F**, middle) indicating the very first moment when the animal starts receiving new information that one wall is suddenly fast approaching.

Decision execution time is estimated from analysis of videography data (**Fig.2G**) using two additional side cameras (Cam2 and Cam3 in **Fig.1A**) that reveal moments when the fore (RF and LF) and hind (RH and LH) legs are touching the ground (**Fig.2H**, bottom, **SI Video 1**). Characteristic coordinated sequence of diagonal LH-RF and RH-LF strides observed corresponds to trotting, the natural gait animal produces^40^ to accommodate speeds faster than 20cm/s. Individual strides of limbs are also detected as periodic oscillations^28^ in lateral acceleration (**Fig.2H**, middle) with maximum positive (turning left) peak at t=0.29s (**Fig.2H**, middle, red ”3”). Comparison of video frames in Fig.2G with gait reconstruction indicates that left turn is executed by stereotypic push (**Fig.2G**, right, red “3”) provided by RH leg (red “3”in **Fig.2H**, middle). This peak in lateral acceleration (SI Methods) is repetitive for all 414 trials for both left and right turns for all animals (**Fig.S7**) and is used to extract trial reaction time (RT). Corresponding time histograms for FT and RT (**Fig.2I**) indicate the choice period of 0.32s between the mean FT (0.06±0.03s, mean ± s.d.) and mean RT (0.38±0.07s, mean ± s.d.).

### Stages of decision-making strongly correlate with epochs of neural activity modulation

During epoch 1 both contra- and ipsi-lateral whiskers periodically palpate walls (**Fig.2F**, middle, black bars) providing continuous sensory information to enable CL wall tracking^29^. Similarly, during epoch 5, periodically varying sensory inputs are generated by oscillating whisker curvature (**Fig.2F**, bottom). However, for both epochs, a nearly stationary neural activity is observed for both population- and trial-averaged activity (**Fig.1J,K**) as well as for single unit activity for all trials (**Fig.S5**) that indicates strong adaptation^18^ of neural population to these repetitive palpating events. While stable during these epochs, the population activity exhibits high variability at ms-scale.

Identifying contribution of behavioral and sensory stimulation variables to the variance of neural activity during epoch1, we used multi-parameter generalized linear-nonlinear Poisson model^41^ (GLM) analysis (SI Methods, **Fig.S12**). For some units whisking strongly dominates the tuning (e.g. Layer 4 excitatory unit, top, **Fig.2J**). Other units (e.g. bottom, **Fig.2J**), however, show mixed representations of whisking and locomotion. In fact, multidimensional tuning is characteristic for most neurons (over 80% of units, **Fig.2K**) with a mixture of whisking kinetics (i.e. angle, amplitude, phase), kinematics (i.e. curvature and derivative), and rich representations of behavioral variables (lateral/ forward speed) and gait (legs positions/ acceleration).

Such mixed high-dimensional representations were observed previously in spontaneous brain-wide activity without external sensory input, indicating sensory-motor integration as early as primary visual cortex^42^. In our case, fusion of sensory (whisker related) and behavioral (motor-related) information encoding in the barrel cortex may help active whisker-guided navigation that requires continuous inference^29^ of relative positions of the obstacle (approaching wall sensed by whisking) and mouse body (proprioception) to estimate time to collision. Unsurprisingly, among the strongest correlations is the feedback signal V_wall_ (feedback in Fig,2J,K) that drives the walls and is dictated by animal navigation avoiding collision.

In contrast to adapted stable spiking during epochs 1 and 5, a single contra-lateral whisker touches an approaching wall at FT (**Fig.2E**, middle, t=0.07s) mirrors a strong increase of neural activity during epoch 2 (**Fig.2K**). Note that a similar peak (peak 2’ in **Fig.1O**) is also observed at the end of trial at t=1.84s, appearing in both population-averaged activity (**Fig.1O**) and single unit response (**Fig.S5**). Here, at the trial end, clamped whisker is released, returning to full range of whisking when the right wall starts to retract (**Fig.S8A**). Therefore, epochs 2 and 2’ are likely associated with coding of mismatch signal^43^ notifying the downstream circuits on the appearance of a sudden deviance^44^ in sensory flow. This is supported by observation of similar peaks during right turns (**Fig.1O, Fig.S5**) that match FT of ipsi-lateral whisker last touch of retracting (t=0.06s, epoch 2) and first touch of approaching (2.15s, epoch 2’) left wall. In all cases, epoch 2 peaks appear within 10ms range from FT consistent with functional afferent delay from whisker follicle to barrel cortex^45^.

Peak 4 activity is separated from peak 2 by a large suppression dip in epoch 3 and is delayed by almost 200ms relative to FT, exceeding possible afferent delays. Peak 4 activity ramps up and reaches maximum while periodic wall-palpating events continue even when the wall starts to approach (mean 2-3 wall touches, **Fig.2F**, middle) until the whisker is clamped at t=0.2s. Hence, it seems unlikely that activity ramping during nearly stationary sensory input can be attributed directly to purely sensory modulation. To further identify the origin of peak 4, we analyzed experiments where mouse runs intermittently with brief stops (1-2sec) during otherwise continuous running at high speeds (**Fig.2L**). While peak 2 is dominant for both ipsi- (**Fig.2L**, blue) and contra-lateral (**Fig.2L**, black) turns when the animal continues active whisking during brief stops, peak 4 activity dominates the modulation only when they are actively navigating (**Fig.2L**, red). Hence peak 4 is likely to reflect participation of primary somatosensory cortex in motor preparatory activity, often observed in higher motor cortices^12–14^. Indeed, peak 4 starts after FT (**Fig.2I**, blue), reaches maximum at the start of RT (**Fig.2I**, red), and is quenched when the decision is fully executed, despite stimulus signals continuing to arrive (**Fig.2F**). Analysis of intrinsic dimensionality^46^ (SI Methods) during decision making (**Fig.2M**), high-dimensional neural coding during epoch 1 collapses rapidly to just a single dimension following the FT during epochs 2, 3 and 4. Using GLM (examples in Fig.S12BC), the majority of units show no significant tuning for both sensory and behavioral variables during decision that they previously exhibited during epoch 1.

Decline of spiking variability at sensory onset is a known universal cortical phenomenon assigned to stabilization of a network state^47^. However, in our experiments the neural population remains one-dimensional^48^ during choice period between FT and RT and starts to increase only at onset of decision execution and epoch 5. Emergence of synchronous modulation^36,37^, dimensionality collapse^48^, and strong suppression locked to sensory stimulation^49,50^ are known neural correlates of top-down regulated attention^38^ and decision making processes^3–5^. In what follows we argue the neural activity ramp during epoch 4 is consistent with a bounded integration mechanism of decision making^3–5,7^.

### Could neural activity modulation be explained by simple sensory drive?

Before diving into argumentation based on decision making interpretation, we first focus on an alternative hypothesis that signals may simply be sensory driven: the wall movement corresponds to variation of whisker stimulation, reflected in corresponding modulation of neural activity. To test and falsify this hypothesis, we performed detailed analysis of correlation of trial-by-trial whisker contact times (**Fig.3B,C**) to epochs of neural activity modulation (single L4 unit activity **Fig.3D, E**), wall movements (**Fig.3A**), and animal strides (**Fig.3F,G**).

**Figure 3.**
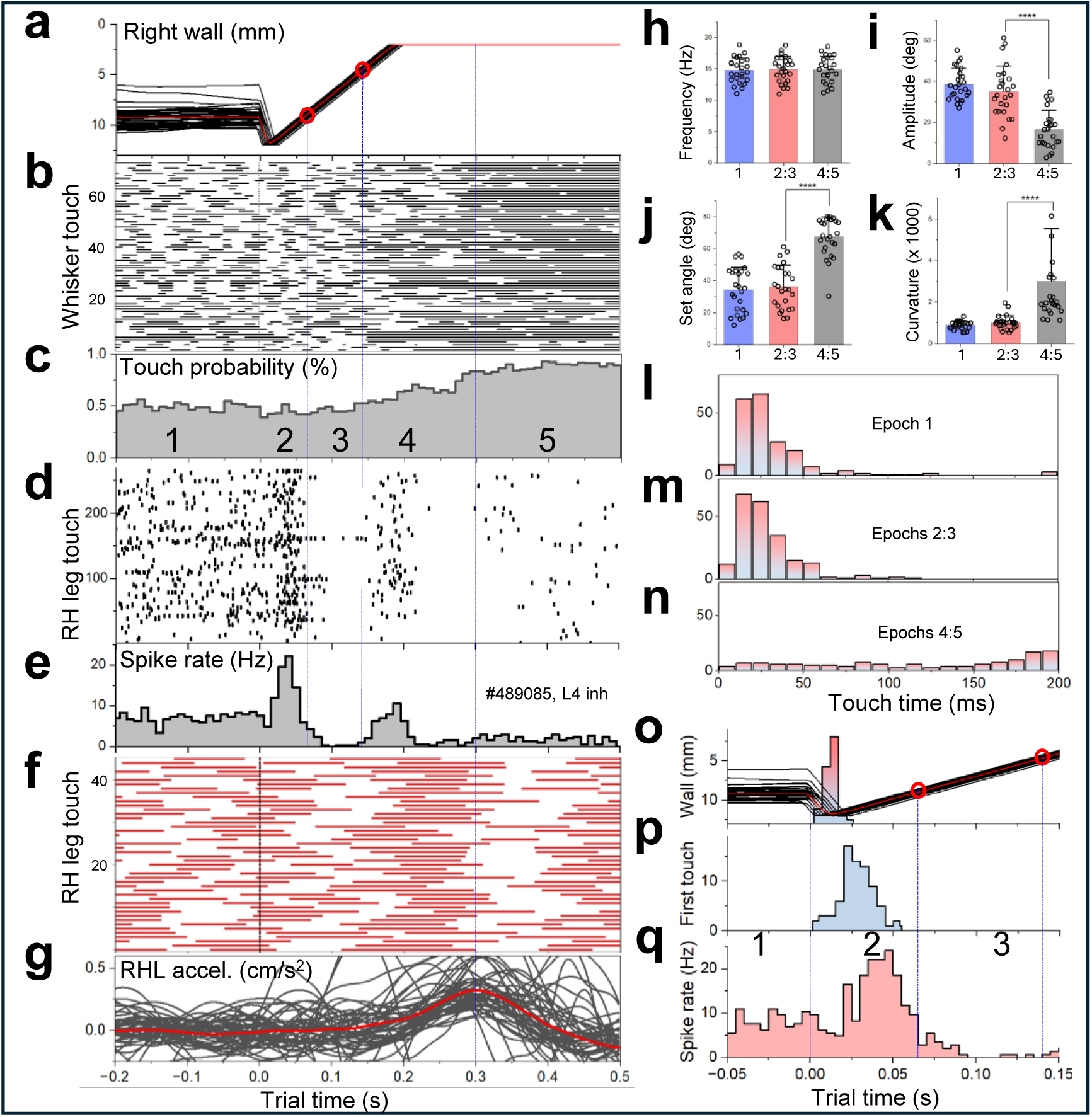
Neural activity modulation is not defined by simple sensory drive. **A)** Wall trace from 50 OL_L turn trials. Red corresponds to mean. First vertical line corresponds to trial switch (end of epoch 1). Second line corresponds to beginning of suppression in epoch 3. Third line corresponds to beginning of ramping in epoch 4. Final line corresponds to beginning of epoch 5. **B)** Raster plot of whisker-wall contacts for 75 trials. Black bars indicate whisker is touching the wall for that time frame on trial-by-trial basis. Extracted from high fps whisker tracking **C)** Whisker-wall touch probability averaged across the same 75 trials as B). **D)** Spike raster plot for example single layer 4 excitatory unit from the same segment across 250 trials. **E)** Peristimulus time histogram (PSTH) for the same unit as D). Using the same trials. **F)** Raster plot of right hind limb strides across 50 trials. Red bars indicate times when animals’ right hind limb is in contact with the ball **G)** Lateral acceleration traces across the same 50 trials in F). Red line indicates mean. **H)** Whisking frequency statistics for example 25 trials across neural epochs, separated into epoch 1, 2&3 and 4&5. Bar shows mean + standard deviation. **I)** Whisking amplitude statistics for same 25 trials as H) across same split of neural epochs, Bar shows mean + standard deviation. Two sample t test p value < 0.0001. **J)** Whisking set angle statistics for same 25 trials as H) across same neural epochs, Bar shows mean + standard deviation. Two sample t test p value < 0.0001. **K)** Whisker curvature statistics for same 25 trials as H) across same neural epochs, Bar shows mean + standard deviation. Two sample t test p value < 0.0001. **L)** Distribution of whisker-wall contact durations using same 75 trials as in B) & C) for epoch 1. **M)** Distribution of whisker-wall contact durations using same 75 trials as in B) & C) for epochs 2 and 3. **N)** Distribution of whisker-wall contact durations using same 75 trials as in B) & C) for epochs 4 and 5. **O)** Overlaid wall trajectory traces for 50 trials, with red line showing mean. Distribution overlaid shows the time point the wall is at maximum distance for each trial (mean = 11ms). Red dots and blue lines correspond to same time points as in A). **P)** Distribution of first whisker-wall touch times after trial switch from the same trials as in O) **Q)** Spike rate PSTH for example unit over the same 50 trials as in O) & P).

To demonstrate that rapid removal of the wall could not result in a decrease of sensory input (hence appearance of suppression dip during epoch 3), we present a zoomed-in view (**Fig.3O,P,Q**) covering epochs 1-3. The wall retracts rapidly (**Fig.3O**), reaching maximum 12mm distance at mean delay of 11ms. Therefore, majority (>80%) of first whisker touch (FT) events (**Fig.3P**) correspond to whisker palpating approaching rather than retracting wall. The strong suppression of neural activity in epoch 3 falls in time range [+65ms: +140ms], corresponding to second touch event, when wall is fast approaching from 8.7mm to 4.4mm (red dots in **Fig.3A** and **Fig.3O**) within direct reach of the whisker. Therefore, reduced firing rate in epoch 3 could not be explained by decrease in sensory stimulation.

During epochs 2-3 wall continues approaching the snout (**Fig.3A,O**) at constant speed until the 2mm position is reached at 200ms. Whisker palpates the wall at constant 15±2Hz frequency (**Fig3H**), with statistically identical amplitudes (**Fig.3I**), and set angles (**Fig.3J**). Itr stays in direct contact with the wall about 48±4% of the time (**Fig.3C**) with median touch duration of 21ms (**Fig.3L,M**), producing periodically modulated curvature with statistically identical maximum per cycle (**Fig.3K**). Whisker touch probability (**Fig.3C**) increases monotonously during epoch 4, reaching 90% in epoch 5. However, while sensory input is monotonous, the neural activity is highly non-stationary with peak in epoch 2, suppression in epoch 3, and another peak in epoch 4 followed by final suppressive epoch 5 when decision is executed by characteristic push by RH leg (**Fig.3F,G**). Therefore, our experimental observations falsify the sensory drive hypothesis and alternative explanations are required.

### Ramping neural activity reflects temporal accumulation of sensory evidence

Canonical framework of perceptual decision making^7^ considers the decision process as the linear integration of evidence over time during sequential noisy sensory sampling until a decision bound is reached. A decision variable (DV) in standard drift-diffusion model (DDM)^3–5,7^ (**Fig.4A**) accumulates sensory evidence (drift), accompanied by stochastic internal or sensory noise (diffusion) that, when reaches either upper (correct trials) or lower (error trials) bound, produces a decision at RT. Fitting classical DDM model (SI Methods) to experimental distribution of RT (**Fig.4B**) produces best fit (Bayesian information criterion, BIC of -664, **Fig.4C**) for non-decision time (NDT) of 0.12s (**Fig.4D**), drift K=2.91, bound B=0.59 (**Fig.4E**), and noise σ=0.25 (**SI Table.2**). Similar parameters are extracted from fitting a “full” DDM model that includes variability of initial conditions X_0_ and NDT. Noticeably, appreciable NDT=0.12s from both DDM and “full” DDM (**Fig.4D**) fits well to 0.12s delay of experimentally observed ramping onset at end of epoch 3 for this example animal (**Fig.4B**, bottom, up red arrow). DDM is suboptimal when evidence strength varies, when whiskers periodically palpate the surface. To accommodate this, we introduced additional model components (Generalized DDM model^51^) including exponentially collapsing bounds^6^ and a leaky integrator^30^ that produce a better fit (BIC of -705). Normalized drift K/B that defines the time needed to reach the bound in absence of noise and decay is 11.12. This is noticeably close to the linear slope of ramping peak 4 of

**Figure 4.**
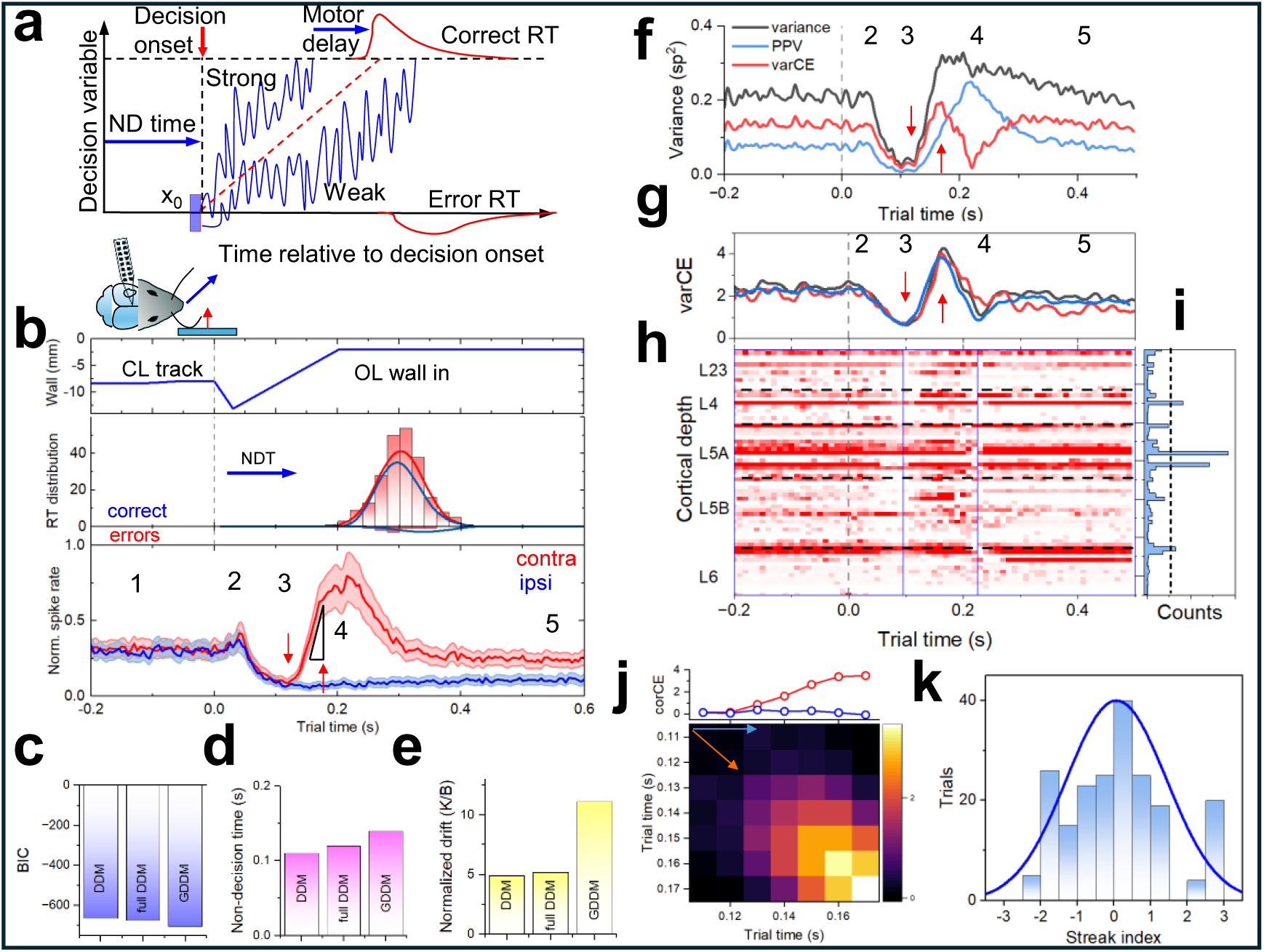
Ramping neural activity reflects temporal accumulation of sensory evidence. **A)** Schematics of drift-diffusion model (DDM) of perceptual decision making. Blue curves are single trial trajectories of a decision variable (DV). The trajectories are affected by constant drift (dashed red line) and subject to stochastic diffusion noise. The decision process starts at delay related to the nondecision time (NDT) and is finished when the DV reaches the upper (correct trials) or lower (error trials) bounds. X0 denotes initial bias in DV. RT are delayed beyond the DV reaching the bound by motor related efferent delay. **B) Top:** Position of right wall during OL_L trials. Middle: RT distribution fitted by DDM (blue) and Generalized DDM (GDDM, red) models. Both produce similar NDT time of about 0.12s shown by blue arrow. **Bottom:** Normalized spike rate for 266 contralateral (contra, red) left and 265 ipsilateral (ipsi, blue) right turn trials. Arrows denote times for beginning and end of the linear ramping activity. Triangle shows linear fitting of the slope. **C)** Bayesian information criterion (BIC) for three fitted models – DDM, full DDM and GDDM. **D)** Non-decision time derived from DDM, full DDM, and GDDM fits to the RT. **E)** Drift normalized on bound derived from DDM, full DDM, and GDDM fits to the RT. **F)** Total variance (black), PPV (blue) and varCE (red) for single unit (489101, RS, layer 5A). Arrows mark start and end of linear ramping activity from B). **G)** varCE variance calculated for all units in same animal (blue), with outliers taken out (red), and calculated on residuals (black). For the same 266 OL_L trials. **H)** Trial averaged activity plot of varCE for same units ordered by cortical depth and assigned to layers. **I)** The sum of varCE calculated for all units in H) for trial time corresponding to choice period (blue lines in H). Dashed line defines 4 outliers. **J)** Covariance matrix CorCE calculated on residuals. Top: corCE values along the matrix diagonal (red) and along the top row (blue). **K)** Streak index distribution for all units across all trials. Blue curve is normal distribution fit.

12.25±0.88 (mean ± s. e. m.) (**Fig.4B**, bottom, triangle) for this example animal (other examples in **Fig.S9** and **Fig.5E** below, n=7, 13.05±4.59, mean ± s.d.). These observations enable hypothesis that onset of ramping activity at 0.12s may indicate the decision onset (**Fig.4B**, bottom, up red arrow), while linear ramping of peak 4 reflects accumulation of evidence until decision is reached (**Fig.4B**, bottom, down red arrow). Thus, a single variable dominating population dimensionality during choice period (**Fig.2M**) may reflect a linearly increasing DV that is reset at the end of epoch 4 when the decision is reached and executed^52^.

**Figure 5.**
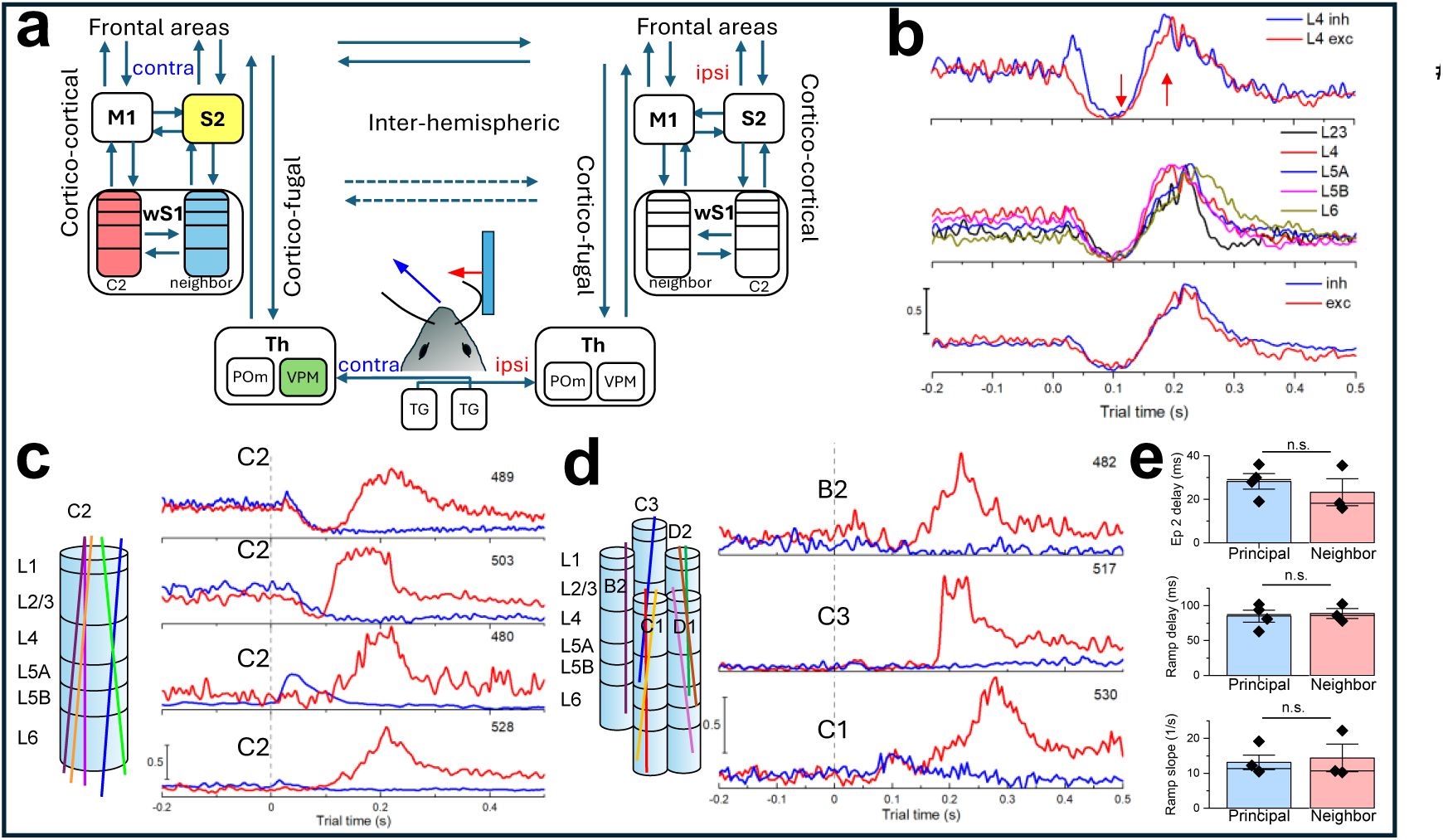
Accumulation of evidence is likely provided by feedback. **A)** Schematics of ascending and descending pathways in whisker somatosensory system. Starts with characteristic isomorphic representation of the whiskers via barrelettes in ipsilateral brainstem trigeminal (TG) nuclei, then barreloids in thalamic (Th) posterior thalamic (POm) and ventroposterior medial thalamic (VPM) nuclei, which project to primary S1 and secondary S2 somatosensory cortices as well as motor areas (M1). Primary (wS1) and secondary (wS1) vibrissae cortical areas have strong reciprocal connections to Th and between themselves. While direct interhemispheric connections exist between contra- and ipsi-lateral S1 and S2, they are stronger in higher order cortices. **B)** Top: Normalized spike rates of subpopulations of FS (blue) and RS (red) units in layer 4. Middle: Same for subpopulations of units in different cortical layers (L2/3 = 10 units, L4 = 13, L5A = 13, L5B = 16, L6 = 13). Bottom: Same for all FS (blue, 43) and RS (red, 22) units. **C)** Left: Schematics of reconstructed probe positions for n=4 animals where recording was in C2 principal whisker barrel (red dots in Fig.1G). Right: Population-averaged rate for these animals with red curves corresponding to OL_L turn trials, while blue curves to OL_R turns. Details for recordings in Table S1. **D)** Left: Schematics of reconstructed probe positions for n=3 animals where recording was done in whisker barrels neighboring C2 (blue dots in Fig.1G). Right: Population-averaged spike rate for these animals with red curves corresponding to OL_L turn trials, while blue curves to OL_R turns. Details for recordings in Table S1. **E)** Statistical comparison of epoch 2 delay (top), delay of ramping activity (middle), and linear regression slope (bottom) from recordings in the principal (left, blue) and neighboring (right, red) barrels. Data represented as mean ± s.e.m.; n.s., not significant (p>0.05), two-sample t-test (top = 0.109, middle = 0.140, bottom = 0.970).

Important signature of perceptual decision-making is gradual accumulation of noise^5,53^ expected for most accumulation-to-bound decision models. Total variance of this doubly stochastics process becomes a sum of two separate components^53^ (SI Methods): point process variance (PPV), proportional to mean spike rates, and variance of conditional expectation (varCE),reflecting variance of rates. For an example unit (**Fig.S10A**) total variance (**Fig.4F**, black) follows PPV (**Fig.4F**, blue) for epochs 1-3, but saturates for epoch 4, when PPV is ramping up. Resulting varCE (**Fig.4F**, red) exhibits fast increase at beginning of epoch 4, indicating accumulation of stochastic trial-by-trial diffusion noise^5,53^. This unit, however, is an outlier with the highest spiking rate and hence abundant noise. When varCE is calculated for each unit in the population (**Fig.4H**) some indeed contribute the most (**Fig.4I**). However, the population-averaged varCE (**Fig.4G**, blue) is similar to that with outliers removed (**Fig.4G**, red). When varCE is calculated from residuals^5,53^ (SI Methods), it is similar (**Fig.4G**, black) showing linear increase between decision onset (**Fig.4G**, up arrow) and decision termination (**Fig.4G**, down arrow) times. VarCE exhibits a fast decline at decision end, reaching minimum at t=0.23 when the decision starts to be executed. Overall, varCE exhibits a linear increase during epoch 4, a characteristic signature of diffusion process consistent with accumulation-to-bound model^5,53^.

It has been argued^53^ that if variance of spike rates during evidence accumulation are defined by a diffusion process, then expectation of spike counts in consecutive time bins (CorCE) should co-vary. The calculated (SI Methods) CorCE matrix (**Fig.4J**) indeed exhibits characteristic increase of correlation as a function of time and, for given time (e.g. along the top row), shows sequential decay as often observed^5,53^ during decision making. This correlation disappears when bins are randomly shuffled (**Fig.S10C**). Together, the time course of CorCE and linear increase of VarCE is indicative of accumulation of evidence during linear ramping in epoch 4. In addition, Streak index analysis^54^ performed on a trial-by-trial basis^55^ (**Fig.S10D**, SI Methods), that shows randomness of local spike count fluctuations with close to zero mean (0.06±1.36, mean ± s.d., **Fig.4K**), indicated gradual rather than step-like transition in each trial during ramping.

## Discussion

Summarizing, during ethologically relevant behavioral paradigm, mice with just C2 whiskers navigate using only sensory information from their whiskers to make high speed navigation-based decisions. We observe consistent stereotyped activity across the cortical column in wS1 that corresponds to the time frame between the first input at FT and decision execution at RT, with 5 distinct epochs. The argumentation presented above falsifies the hypothesis that this modulation is related to simple sensory drive and provides strong arguments towards alternative explanation. The epoch 4 ramping arises at delays considerably longer than sensory afferent delays. Ramping occurs while sensory input stays nearly stationary. Ramping reaches its maximum and starts to decay during onset of motor decision execution at RT while stimulus signals continue to arrive. Epochs 3 and 4 are accompanied by drastic collapse of high-dimensional neural activity to a single latent variable. Fitting of RT distributions to bounded accumulation models (DDM, GDDM) reveals this single variable is consistent with linear evidence accumulation of DV. Fitted drift slopes and non-decision times in behavior data are consistent with observed ramping slope and ramping delay in recorded neural activity. Analysis of spike count noise during this period showed characteristic linear increase in internally driven spike rate variability (varCE) and decrease of its correlations (corCE) consistent with the gradual accumulation models of decision making^53^. Streak index analysis^54,55^ also supports this interpretation. Overall, these 9 arguments provide evidence that epochs in modulated neural activity observed during active single-whisker-guided navigation are likely to represent stages of perceptual decision making.

First-pass prediction of accumulation to bound models is that ramping slope should be decreasing with decrease of stimulus strength. To test this experimentally, we focused on the trial end when the contra-lateral wall moves out (t=1.8s) from being at 1mm proximity to 10mm mean distance during subsequent CL wall tracking (**Fig.1J**, top). Epoch 2’ peak coincides with release of contra-lateral whisker from being strongly bent during the turn (**Fig.S8A**), corresponding to the lower part of psychometric curve (**Fig.2C**). Epoch 3’ suppression is significantly longer with the onset of ramping activity at t=2.0s (0.21s delay) suggesting regulation of integration onset to task demand. Ramping in epoch 4’ is much slower reaching maximum at t=2.58s (0.78s delay). The fitted linear slope (2.05±0.05, mean ± s.e.m., black triangle in **Fig.1O**) is 7X smaller than for the first turn at trial start (12.25±0.76, mean±s.e.m., triangle **Fig.1N**) consistent with drift-diffusion model prediction. Similarly, accumulation to bound models predict decrease of the stimulus strength should be accompanied by accumulation of larger uncertainties in decision making (**Fig.4A**), consistent with observed 2X wider distribution of lateral distances at trial end relative to trial middle (2.30 s.d. at 3 s trial time vs 1.49 s.d. at 2 s time for **Fig.2B**, top, see also **Fig.S6B**, top). Slower ramping is accompanied by longer and broader distribution of angle derivative peaks (**Fig.S8D**) although exact identification of RT is difficult as decision execution is distributed among several consecutive strides.

Where are these computations taking place? Most information about whisker contact with objects is relayed through lemniscal pathway^8,15,45^ (**Fig.5A**). Can ramping signal be computed locally within a principal barrel column in primary wS1 (**Fig.5A**, red)? All epochs including epochs 3 and 4 are already present in thalamo-recipient Layer 4 populations of fast spiking (FS, putative inhibitory, **Fig.5B**, top) and regular spiking (RS, putative excitatory, **Fig.5B**, top) units. They also dominate the activity in sub-populations of all units across different cortical laminae (**Fig.5B**, middle panel) and all excitatory and inhibitory units (**Fig.5B**, bottom panel). Similar synchronous modulation of is observed for other animals (n=4, **Fig.5C**) with recordings from principal barrel column indicating that the coding scheme behind epochs 3 and 4 is unlikely to be produced by local columnar circuitry but rather regulated from outside of the principal whisker column.

Can epoch 3 and 4 modulation involve surround/lateral inhibition^56^ from neighboring barrels columns^57^ (**Fig.5A**, blue)? Cross-columnar excitatory long-range axon collaterals connect neighboring columns running predominantly along barrel rows^58^.

However, staged and synchronous modulation epochs observed in animals where recordings were done from a neighboring barrel in the same or even different row (n=3, **Fig.5D**), exhibits a striking similarity indicating lateral cross-columnar inhibition^57^ is unlikely a source. Indeed, delay of epoch 2 (**Fig.5E**, left), start of ramping at end of epoch 3 (**Fig.5E**, middle) and a linear slope of ramping activity of epoch 4 (**Fig.5E**, right) shows no statistically significant difference (two sample t-test, p>0.05) between principal C2 whisker and neighboring barrel columns. This agrees with conjecture that coding scheme behind epochs 3 and 4 is unlikely to be produced within barrel cortex but rather is a signature of a global feedback regulation.

If epochs 3 and 4 are not generated locally, where can they arise from? Cortico-thalamo-cortical feedback loops^23,59,60^ (**Fig.5A**, green) or initiated downstream in higher order cortices like S2^22^ (**Fig.5A**, yellow) or even hierarchically higher frontal areas and then top-down regulated back into S1?

For observation of remarkable inter-hemispheric asymmetry in contra-lateral (left turns) and ipsi-lateral (right turns) trials (**Fig.5CD**, see also **Fig.1J-O**). While epochs 2 (at trial start) and 2’ (at trial end) are present for both left and right turns, epochs 4 and 4’ are exclusively for contra-lateral left turn trials indicating extraordinarily strong lateralization. Therefore, this global regulation is unlikely originating from frontal cortical pre-motor areas^12–14^ since frontal cortices have a high degree of symmetry for both ipsilateral and contralateral projections^61^. In contrast, sensory S1 and S2 cortices have strongly lateralized ipsilateral outputs^45,61^ and are densely and reciprocally interconnected^62^ forming a top-down feedback loop that exhibits choice-related activity^10,11,22–25^.

Similarly, cortico-fugal^63,64^ and, cortico-thalamo-cortical^59,60^ organization is also mainly ipsilateral and known to form feedback control loops^23^. For example, posterior thalamic nucleus (POm) may provide a thalamic relay for cortico-cortical information flow between S1 and S2 sensory areas^59^ and was shown recently to be indispensable in whisker-based texture discrimination tasks^60^. Therefore, both cortico-cortical and cortico- fugal feedback loops are likely to contribute to observed modulation epoch. Our experiments indicate, however, that contra-lateral activity epoch 4’ continues ramping unperturbed at trial end (**Fig.1O**, red) when ipsi-lateral whisker is touching approaching wall (t=2.16s, 0.36s delay from the onset of wall movement) to generate a strong peak 2’ (**Fig.1O**, blue). Indicating two processes are likely orthogonal to each other, originating from different loops. Our findings are consistent with the hypothesis that “early”^20,21^ sensory response (peak 2 in our case) is mostly perception-related, involving thalamo-cortico-thalamic loops^60^, while “late” mostly decision-related response^19^ (peak 4 in our experiments) reflect reverberation of neural activity^26,30^ in cortico-cortical S1-S2 loops^10,11,22-24^.

A limitation is focusing on analysis of decision-related activity exclusively from primary S1 area. However, our experiments produce intriguing observations pointing to architectural organization of feedback cortico-cortical and cortico-fugal loops contributing to observed decision-related ramping activity in S1. To study transformations of sensory inputs to decision-related variables would require simultaneous recording from multiple areas potentially involved in feedback regulation ^10,11,22–24^. We demonstrated here that these dense neural activity datasets during our single-whisker-guided ethological decision-making behavioral paradigm is a powerful approach for mechanistic dissection of complex nested loops. Therefore, offering a key step toward understanding the contribution of early sensory cortical circuits to decision making during natural behavior.

## Author Contributions

AGA designed research, conducted experiments, analyzed data, and wrote the draft; YV acquired funding, built behavioral rig, designed research, analyzed data, wrote the draft and edited the final text.

## Acknowledgments

YV is a CZ Biohub Investigator. Authors are grateful to Kun Hu and Nur Al-Kodmany for help in setting up the VR rig and development of behavioral protocol and to Ksenia V. for insightful discussions.

## Methods

### Animal preparation

This study is based on data from 19 mice of both sexes aged between P65 and P112 (**Table.S1**). To ease identification of principal barrel and enable optogenetic tagging, the transgenic Scnn1a-TG3-Cre (Jackson:009613) line was crossed with the Ai32 line (Jackson: 012569), producing strong co-expression of the yellow fluorescent protein (YFP) and Channelrhodopsin-2 (ChR2H134R-EYFP) in ∼85% of layer 4 excitatory neurons^1^ in the primary somatosensory cortex (**Fig.1F**).

All procedures were in accordance with protocols approved by the UIUC Institutional Animal Care and Use Committee. Animals were kept on a reverse 12-hour light/dark cycle in individual cages equipped with activity wheels (K3250 Fast-Trac, Bio-Serv) to encourage active behavior in-between experimental sessions. All experimental sessions were performed during the dark phase. Surgical procedures were carried out aseptically while animals are anesthetized with isoflurane (2-4% in oxygen) before being injected subcutaneously with 5mg/kg carprofen. To prepare animals for behavioral and electrophysiological studies a custom-built titanium headbar (10mm x 2 mm)^2^ was attached to the animal’s skull to hold the animal’s head. Orientation of the headbar is carefully adjusted with respect to Bregma and Lambda points on the animal skull to provide a repeatable coordinate system for electrophysiological multielectrode probe insertion. The central open area of the headbar (3mm x 2mm) exposes the skull directly above the primary somatosensory cortex for in-vivo visualization, targeted craniotomy, and precise probe insertion. This area is covered in optical grade clear cement for in-vivo fluorescence imaging. For experiments with electrophysiological recordings, a craniotomy (approximately 0.2 mm in diameter) is made through the back of the headbar, and a ground wire (2 mm long Pt/Ir ground wire, A-M Systems 776000) is implanted in the cerebellum and cemented on the headbar. Beginning the day following the surgery, subcutaneous Carprofen (5mg/kg) is administered for two days to reduce inflammation. To restrict sensory information flow, all but a pair of C2 whiskers are trimmed at the time the animal is placed on water deprivation. Trimming is performed daily afterwards to avoid other whiskers growing.

Following full and complete recovery from a previous surgery, the animal is placed on a water restriction schedule^2^ in preparation for behavioral conditioning. Food is continuously available. Water is adjusted (approximately 1 ml per day) so that mice maintain more than 80% of free-drinking weight. When fully adjusted to the water restriction protocol (weight stabilized) a mouse is habituated to virtual reality treadmill of **Fig.1A** with head fixation starting with 15min during the first day. Using a large diameter sphere (16 inches) in our VR rig as well as tunable head fixation enables to closely mimic the posture observed during natural locomotion^3^. Sessions are extended over 3-4 days to last up to 1 hour. During this time a mouse is encouraged to run by administering water rewards (0.1ml) given only if the running speed exceeds 5cm/sec for a total of 1 ml. Typically (**Table S1**), on day 3 a mouse is fully acclimatized to the apparatus, actively runs 80% of the time at elevated speed above 10cm/s (21±4cm/s, mean ± s.d., **Fig.S1A**) and consumes all the daily water ration during experiment. When running, they navigate the straight VR corridor using only a pair of C2 whiskers while minimizing the lateral displacement to less than ±1cm (**Fig.S1B**).

### Behavioral tactile virtual reality rig

The treadmill of **Fig.1A** is built around a hollow Styrofoam sphere (16-inches diameter) with walls custom-thinned to provide a total weight of less than 60g. The sphere is suspended on 7 ping-pong balls supported by 7 air-nozzles distributed around its bottom half^4–7^ based on the design developed at the HHMI Janelia Research campus^8^. Controlled stimulus to animal whiskers is provided by a pair of motorized walls positioned at both sides of the snout. The animal movement in lateral (Vlat) and forward (Vfor) directions is tracked at 500Hz by recording the sphere movement with two tracking cameras^9^. Turns were defined by a turn angle 𝜃 that was positive for left turns and negative for right.

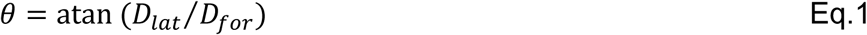

where 𝐷_𝑓𝑜𝑟_ and 𝐷_𝑙𝑎𝑡_ are lateral and forward displacements in a single cycle. This signal is used to provide positive feedback to the movement of the walls (Vwall):

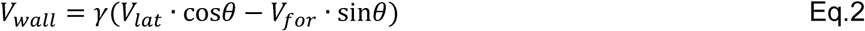

where 𝛾 is a calibrated feedback coefficient. When taken as negative it generates a positive closed-loop (CL) virtual reality (VR) that moves walls closer to the snout if the animal is moving closer to them and, therefore, encourages the animal to stay in the middle of the VR corridor.

The real time (RT) control and acquisition system is custom designed to power a FPGA CompactRIO (cRIO-9045, NI) with 32 channel analog voltage input module (NI 9205) to read sphere velocities, 4 channel analog output module (NI-9263) to operate mechanical walls, and 32 channel TTL input/output (NI-9403) to provide synchronization to video cameras and other equipment. The module is operated via LabVIEW RT Module (NI) by the custom designed LabVIEW acquisition and control program^7^ that has Master/Slave architecture. The program operates several parallel processes that need to run simultaneously and continuously but at different rates. The master process runs in a loop at 30MHz and provides RT analog acquisition and control data to the rig. The master loop controls several slave loops and communicates with them using messaging architectures. One of the slave loops buffers recorded data and logs the data to disk at its own much slower pace. As a result, the acquisition and control experimental protocol executed in the master loop can run continuously without interruptions with virtually no inter-trial interval. Another slave loop is designed to provide a graphical user interface (GUI) that allows the user to observe data on the screen such as overlaid trial trajectories of the animal and to change parameters for each trial session if needed. Control signals generated in the master loop are directed to synchronize 3 high-speed cameras (**Fig.1A**) to acquire videos of whiskers and paws, as well as to synchronize behavioral recordings with electrophysiological recordings, optogenetic stimulation, and provide water rewards to the motorized licking port if needed.

The VR rig is enclosed in a light-tight box with fine copper mesh on the inside to provide shielding from ambient light and from EM interference. Constant hissing sound of compressed air from the air nozzles on which the VR sphere is suspended on (79.9±0.2dB SPL measured with calibrated microphone over 1-100KHz range) masks all background sounds including <25dB noise from motors moving the walls. The rig itself is positioned in a fully sealed light-attenuating box, with the only illumination used for cameras being via infrared LEDs, which use IR wavelengths well beyond the limits of mouse visual system. Additionally, during behavior experiments, all lights in the room are switched off and the box is covered in additional laser safety curtain. The surface of the Styrofoam sphere is coated with permanent water-resistive protective coating using low-odor latex aerosol (Krylon 7120) to enable the sphere to be cleaned between each animal thus minimizing olfactory cues.

Therefore, the only sensory cues left for animals to use for navigation in the VR are whiskers palpating the motorized walls.

### Stereotaxic-less probe positioning on a principal whisker barrel

We aim for precise placement of microelectrode probe strategically into the principal whisker barrel and to align it perpendicular to cortical layers. We also aim to produce active behavior when the head-fixed animal is running fast on a treadmill navigating virtual turns for the whole duration of the electrophysiological recordings. This precludes the use of stereotaxic surgery for electrode insertion. Therefore, we developed a protocol for highly precise probe insertion when the animal is placed, and head fixed in the VR rig.

The first step is to acquire a brain map of the somatosensory cortex generalized across many animals (**Fig.S3D**). For that we used fluorescence top-down imaging (Olympus, MVX10) of the whole fixed brain harvested immediately after the final terminal experiment (**Fig.S3C**). To match vasculature patterns with those observed in-vivo in the same animal (**Fig.S3B**), mice were deeply anesthetized with 5% isoflurane then perfused with 0.1 M sodium phosphate buffer for just a short period of time to ensure that some blood remains in top arterioles on the brain surface. Resulting brain preparations not only show characteristic contours of primary visual cortex (V1), the barrel cortex (wS1), as well as characteristic shapes of lower jaw (LJ) and hindpaw (HP) somatosensory areas (**Fig.S3C**), but also in some cases show identifiable locations of main barrels across main B, C, and D rows (**Fig.1G**). Combining similar images from several animals, a generalized map is created with positions of individual barrels relative to the contours of V1 and S1 areas (**Fig.S3D**).

Next, for a given mouse, to map in-vivo the location of the target C2 barrel without the use of stereotaxic apparatus, the mouse is anesthetized with isoflurane (2-4% in oxygen) and the brain is imaged in-vivo (**Fig.S3A**) through the central open area of the headbar that exposes the skull directly above the primary somatosensory cortex (nominal coordinates for C2 barrel 2.0mm AP, 3.4mm ML). The YFP expression in L4 excitatory cells in the primary somatosensory and visual cortices is strong enough to identify contours of V1, wS1, LJ and HP areas as well as characteristic vasculature patterns through the skull (**Fig.1F, Fig.S3B**), and to align them to previously acquired cortical map (**Fig.S3D**). This procedure allows us to mark the location of the C2 barrel with respect to the headbar with localization precision as high as ±0.1mm as confirmed by subsequent histological mapping (see below) (**Fig.1H**).

### Targeted Craniotomy

On the morning of the final terminal experimental session, mice are briefly anesthetized with isoflurane (2-4% in oxygen) and are placed in a head-fixation surgery apparatus closely mimicking head-fixation setup in the VR rig. Fluorescence imaging is performed with overhead stereo microscope (MZ12.5, Leica) equipped with fluorescence adaptor (Quad Fluorescence System, Kramer Scientific). Brain map with location of targeted C2 principal whisker barrel generated during in-vivo fluorescent imaging (**Fig.S3B, D**) is compared with microscope imaging to check the marked location of targeted craniotomy on the skull surface. Using a microdrill with 0.002” burrs (Fine Science Tools, Canada), the optical cement and the skull are carefully thinned while the surface is cooled down using sterile artificial cerebrospinal fluid (ACSF). A small craniotomy of < 300µm diameter is opened while the dura remained intact. The craniotomy site is then imaged both in a white light and with fluorescence filters to visualize a barrel field and a vasculature pattern and to produce a final map for probe implantation. The craniotomy site is covered with a silicone elastomer plug (World Precision Instruments, USA). The whole procedure typically lasts less than 30 mins and mice are returned to home-cage for recovery for at least 2 hours. Typically, mice are fully recovered within 20 mins and frequently running on activity wheels.

### Electrophysiological recordings

For extracellular electrophysiological recordings we used either (**Table S1**) a 64-channel linear shank silicon probes (Cambridge NeuroTech, UK) connected to Intan head stage (Intan, USA) or Neuropixels (1.0, Imec) electrode arrays with 383 recoding sites selectable from a total of 960, connected to custom PXIe acquisition module (Neuropixels, Imec). The acquisition board (OEPS-9030, OpenEphys) and Open Ephys GUI were used to demultiplex and visualize recorded voltages. The TTL signals from the RT LabVIEW acquisition program were used to synchronize recordings in real time with behavioral sessions. Probes were mounted to a custom 3D-printed holder and attached to a steel rod held by a robotic micromanipulator (MPM System, New Scale Technologies) at 38 degrees around the animal head AP axis to ensure shank was perpendicular to cortical layers in the C2 barrel column. The inner surface of the probes was painted with DiI (V22888, ThermoFisher) for post hoc probe track visualization during histology.

Once mice are placed and head-fixed in the VR rig they almost immediately start to run. To direct animals, the behavioral session starts right away with closed loop CL straight run. The animal skull with a craniotomy site mapped previously is observed with overhead stereo microscope (MZ12.5, Leica), plug is removed, and a silicon probe is positioned over the C2 principal whisker barrel and slowly implanted at constant speed (50-200μm/min) to a full cortical depth of 1mm. Once inserted, the probe was left to settle for at least 5 minutes prior to the start of the main behavioral session. For most sessions the craniotomy site was covered with a saline sterile ACSF bath. In some experiments saline-based agar was used covered with silicone oil to prevent drying.

### Optogenetic and histological verification of electrode position

At the end of every electrophysiological session an in-vivo optogenetic stimulation of layer 4 was performed using fiber-pigtailed 473nm laser light (10 Hz with 2 ms pulses, 1 mW peak) focused on the craniotomy area to obtain current source density (CSD) traces. Characteristic negative sink at short time scale after the optical pulse is used to identify the position of layer 4 (**Fig.1I**).

In sessions where 64-channels probes were used (**Table S1**), small electrolytic lesions were made using the lowest electrode of the silicon probe to ease identification of the electrode depth in histology (arrow in **Fig.1H** and **Fig.S3E, F**). For that purpose, corresponding pins in a head stage board were connected via external wire to a constant current isolated stimulator (DS3, Digitimer). Then 10µA of current was sent through the wire to the lesion electrode for 500 ms while the head stage was disconnected.

Immediately after the end of final terminal experiment, mice were deeply anesthetized with 5% isoflurane then partially perfused to enable vasculature to be visible. Harvested brains removed from the skull were immediately imaged with top-down fluorescence microscopy that enables matching of postmortem vasculature patterns (**Fig.1G** and **Fig.S3C**) with those observed in-vivo (**Fig.1F** and **Fig.S3B**) and to identify the DiI-marked entry point of the probe (**Fig.1G**) with the barrel field map (**Fig.S3D**) using image overlay alignment (Photoshop, Adobe Suite). The brains were then sectioned into 70-μm-thick coronal sections (VT1200S, Leica) to allow for visualization of DiI-stained probe tracks, electrolytic lesions (**Fig.S3E**), and identification of cortical layers (**Fig.S3F**). Strong YFP expression in Layer 4 clearly identifies individual barrels (**Fig.S3F**). Dense projections to layer 2/3 terminate at the border with layer 1 that ease its identification. Layer 6 has the lowest YFP intensity (**Fig.S3G**) enabling further identification of the border between layer 5 and layer 6. Expression is not uniform across layer 5 with stronger expression at its bottom that is assigned to the boundary between layer 5A and 5B (**Fig.S3G**). Combination of YFP intensity distribution, lesion location, the location of DiI probe traces, as well as the CSD analysis were critical for localization of the laminar position of the recording electrodes (**Fig.1G**). Each electrode was then assigned a depth value that is later used to assign recorded units to cortical depths and layers within a given barrel column (**Fig.5C, D**).

We also registered electrodes and units to the Allen Institute Common Coordinate Framework (CCFv3)^10^ by marking 20 registration points in consecutive coronal slices. Fluorescent probe tracks were manually labelled and used to translate the probe track positions into the CCFv3 coordinate space. We found, however, that manual reconstruction of electrode positions with respect to positions of L4 barrels and cortical layers individually for each animal using methodology described above was more precise than provided by a global CCFv3 mapping that does not define individual barrels. Therefore for our analysis we used our manual reconstructions of a probe position for the principal and neighbor barrels as shown in **Fig.5C-D**.

### Spike sorting and unit filtering

The pipeline for spike sorting and filtering is shown on **Fig.S2A**. The recorded electrophysiology data were automatically spike sorted using Kilosort3 (https://github.com/cortex-lab/Kilosort). The sorted datasets were passed through the Ecephys code for spike sorting pipeline (Allen Institute for Brain Science, Ecephys package, https://github.com/AllenInstitute/ecephys_spike_sorting) using functions kilosort_postprocessing, mean_waveforms and quality_metrics. Quality metrics to filter out putative single units (**Fig.S2B**) were defined by Ecephys default values as amplitude cutoff ≤0.1mV, ISI violations ≤0.5, and presence ratio ≥0.90. We also used waveform amplitude, signal to noise ratio, waveform duration, waveform spread, isolation distance, nearest neighbor’s hit rate, and mean firing rate to further filter units. Finally, the waveforms were manually curated with the Phy GUI (https://github.com/kwikteam/phy). During manual curation, each unit was inspected and units with artefactual waveforms and putative double-counted spikes were discarded. The finalized 634 units (**Fig.S2C**) were classified as fast spiking (FS, putative inhibitory) and regular spiking (RS, putative excitatory) based on their waveform duration using 0.4ms as a threshold (**Fig.S2D**). Electrode position mapping for individual animals enables the assignment of units to cortical depth based on the location of electrode with maximum voltage and to cortical layers (**Fig.S2E**). Distribution of all units across layers (**Fig.S2F**) reveals a typical bias towards layers 5A and 5B with less contribution from layer 2/3, mostly dictated by their sparser spiking and lower spike amplitudes.

Spiking data for all filtered units for each animal for the final terminal session are packaged together with behavioral data into Neurodata Without Borders files for further analysis (NWB2.0, https://github.com/NeurodataWithoutBorders/nwb-schema).

### Whisker tracking and reconstruction of whisker angle and curvature

High speed overhead video of the whisker (**Fig.2E**, **SI Video 1**) was captured simultaneously with the electrophysiology recordings using the overhead (cam1 in **Fig.1A**) 656 × 600 pixels camera at 1000–1000 fps (EoSens1, Mikrotron) equipped with a 0.36× telecentric lens that produced 25 mm × 23 mm field-of-view (Edmund Optics, no. 58-257). Video streams were digitized with a frame grabber (BitFlow Axion) controlled by StreamPix7 multicamera software (NorPix). To align videoframes to the master clock with less than 1 ms jitter, a video system was triggered from the main RT LabVIEW acquisition program. The field of view (FOV) is adjusted (MC ControlTool, Mikrotron) to capture the full length of the C2 whisker as well as its bent shape when interacting with a wall. The whisker was illuminated by an overhead LED light source (ThorLabs M810L3) powered by DC power supply (Mean Well RS-15-24) and focused with 40 mm focal length aspheric condenser (Thorlabs, ACL5040U). Whisker movement during interaction with the wall was tracked by extracting individual lossless frames for subsequent manual analysis or using automated tracking with SLEAP^11^ model (**Fig.2F**). A skeleton of four points along the C2 whisker was manually identified for 50-100 frames for each video (**Fig.S11A,B**) and used as the training dataset for the model. Once trained, the model was used to make inferences on all frames in the video using the Hungarian matching method. The angle of the whisker is defined as 0° when it is perpendicular to the mouse head AP axis. The first two tracking points along the whisker length in each frame were used to calculate the whisker angle.

Curvatures are calculated from fitting a circle to the three points, either closest to the base (points 1, 2 and 3 in **Fig.S11B**) or closest to the whisker tip (points 2, 3 and 4 in **Fig.S11B**). Comparison between base and tip curvatures is shown for an example period of CL straight run (**Fig.S11C**) and OL turn (**Fig.S11D**). The tip and base parts of the whisker are moving in-phase (Fig.S11E,F) with phase shift of 0.2±2.4° (mean ± s.d.) over 100 example cycles. These results demonstrate that higher order modes^12^ are not excited neither in CL nor in OL periods.

### Tracking of fore paws and hind paws and reconstruction of gait

Videos of hind paws and fore paws (**Fig.2G, SI Video1**) were captured using a pair of 659 × 494 pixels cameras at 50fps (Basler acA640) (cam2 and cam3 in **Fig.1A**) equipped with 16 mm lenses that produced 34 mm × 23 mm FOV. Cameras were positioned above the hind limbs and to the side of the front limbs and were triggered from the main RT LabVIEW acquisition program. The FOVs were illuminated by additional LED light sources. Frame-by-frame manual inspection of the paw’s videos (**SI Video1**) enabled reconstruction of the animal gait (**Fig.2H**, bottom) by recording the precise timings of when each paw is in contact with the sphere (**Fig.2G**). This reconstruction is confirmed by observation of characteristic periodic oscillations (**Fig.2H**, middle) in a derivative of the recorded traces of turn angle 𝜃 (or lateral acceleration). Characteristic coordinated sequence of diagonal LH-RF and RH-LF strides corresponds to trotting, the natural gait that animal is producing^3^ to accommodate speeds faster than 20cm/s.

### Data analysis

Data are publicly available online at Dandi: https://dandiarchive.org/dandiset/001341

### Spike rates for individual units and populations

To visualize firing rates for individual units, the spiking activity was binned at 0.005s and plotted as raster plots (**Fig.S5A-B** and **Fig.S10**, bottom) or averaged across all corresponding trials (**Fig.S5C-D** and **Fig.S10**, middle). The raster plots of population activity (**Fig.1J-K** and **Fig.S4**) are combined from individual units z-scored trial-averaged spike rates. Population spike rates plots (**Fig.1L-K** and **Fig.S4**) represent population-averaged spike rates normalized on the number of units recorded for a given animal. For comparison with DDM results (**Fig.4B** and **Fig.S9**) or between animals (**Fig.5B-D**), the population spike rates are presented as normalized values with minimum and maximum spike rates normalized to 0 and 1, respectively.

### Multiparameter GLM regression

To assess which variables contribute the most to the neural activity during closed-loop baseline straight running, we used a multi-parameter generalized linear-nonlinear Poisson encoding model^13^ (GLM). Python code was used available publicly from https://github.com/pillowlab/GLMspiketraintutorial_python.

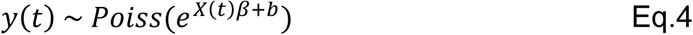

Concatenated spike trains during closed loop CL straight running trials were created for each unit individually. The same 3 second segment was used for 40 randomly selected CL trials. Spikes were arranged into 20ms time bins and smoothed using a Savitzky-Golay filter. Spike counts (y(t)) were predicted using the model with design matrix (X(t)) which included 13 different types of task-related, sensory and movement variables. Model was validated and tested by splitting the data and using five-fold cross validation with weights (β) fit by maximizing likelihood with ridge regularization for each fold. Spike triggered average (STA) was computed and visualized for each of the matrices to produce an unbiased estimator of the filter to be used and determine if any issues were encountered. We then computed a whitened STA; the maximum-likelihood estimator. Pre-whitening is a common method used to remove autocorrelations from residuals before GLM analysis of neural data. The whitened STA was used for model fitting using the ‘glmfit’ python function with Poisson noise model and exponential link function. Spike predictions and subsequent quantification of performance were recorded for each regressor.

For regressions we used a set of simultaneously recorded parameters of whisking, gait, speed and direction. To correlate whisking motor actions that are known to strongly contribute to spiking activity^14–16^, whisking phases and amplitudes were extracted from the tracked whisking angle using Hilbert transform. To correlate locomotion parameters, run angle derivative and times for RH and LH legs pushes were used. Together with speed parameters (forward and absolute speeds) as well as direction parameters (run angle and lateral speed) the pool accounted for 13 regressors. Although they do not form orthogonal basis as some of them depend on each other, however they cover different aspects of sensory input (whisking), locomotion (speed, gait) and motor action output (direction) (**Fig.2J**). Additionally, we included a feedback parameter 𝑉_𝑤𝑎𝑙𝑙_ defined by Eq.2 that appears to produce the highest contribution.

For each individual unit, the log likelihood was calculated for the multiparameter GLM (full model), then one by one each of the variables were removed, with log likelihood calculated on the reduced model. This was done across all 5 splits. Log likelihood ratio was then calculated for each parameter with the average taken across the 5 splits to obtain the test statistics. This statistics was used to obtain a p-value from chi-squared distribution with 1 degree of freedom, as per likelihood ratio test. If the reduced model showed significant likelihood ratio test, then the neuron was assigned as encoding that variable (**Fig.2JK, Fig.S12**)

### Dimensionality reduction

Analysis of intrinsic dimensionality of population activity is based on calculating principal components (PC) on the dataset and estimating the 90% of the data variance that can be explained by a minimal number of PC. The dimensionality reduction code following a standard PCA reduction method^17,18^ is rewritten in Python and is used for analysis in **Fig.2I**. The spiking activity for each unit was taken in 100ms epochs across all trials of the same type, with mean rates calculated in 1ms time bins for each unit. A 2ms gaussian smoothing kernel was used and units with a firing rate <1Hz were removed. Subsequently, for each 100ms segment, a matrix of trial average spike rates was produced and used for PC analysis. The number of latent variables required to explain 90% of the variance was assigned as the dimensionality for that segment.

### Fitting of drift diffusion models

Reaction times (RT) are extracted from lateral acceleration (or run angle derivative) as the largest peak in the time interval after the start of the turn trial (**Fig.2H**, middle). Such peaks are associated with characteristic push with RH or LH legs as observed from videography of the animal locomotion. These peaks are highly reproducible for hundreds of randomized left and right turn trials for all animals (**Fig.S7** and **Fig.S8**). We assigned a correct score to the turn if animal is achieving ±1cm clearance at t=1s from the trial start, otherwise the trial is marked as error. For an example animal in **Fig.2A** the success rate is reaching 95% (14 failed trajectories in red). Similar results are obtained for other animals (**Fig.S6**). Corresponding distributions of correct and error RT are shown in **Fig.4B** and **Fig.S9**. To fit the RT with different drift diffusion models we used a Python based pyDDM generalized drift diffusion model^19^ (https://github.com/mwshinn/PyDDM).

### Analysis of varCE

To calculate varCE and corCE we followed the method^20^. For each 10ms time bin the mean number of spikes for each unit 𝑁̅ was calculated together with the total variance of the spike counts 𝑠^2^ across all left turn trials. The total variance of this doubly stochastics process becomes a sum of two separate components^20^: point process variance (PPV) that is proportional to mean spike rates and variance of conditional expectation (varCE) that reflects the variance of rates.

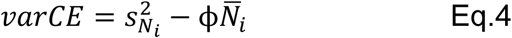

where the constant ϕ was estimated using the time bin that contains the lowest Fano factor that ensures the VarCE remains positive throughout the choice period. The PPV was defined as:

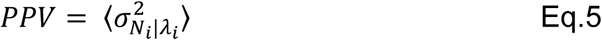

where 𝑁_𝑖_ is the number of spikes in the bin and 𝜆_𝑖_ the firing rate.

For an example unit the total variance (**Fig.S10A**, middle, red) follows the mean (**Fig.S10A**, middle, blue) for epochs 1 and 2, that results in a Fano factor (**Fig.S10A**, top, black) of 0.74±0.04 (mean ± s.d.). Total variance saturates midway through a strong increase of mean firing rate during ramping that results in a Fano factor dropping down to 0.29 at the peak of epoch 4 at t=0.23s and then slowly restores back in epoch 5. If all the total variance at t=0.23s is taken as coming from the PPV, then constant ϕ can be taken as 0.29. The resulting varCE (**Fig.4F**, red) exhibits a fast increase at the beginning of epoch 4 between t=0.12s and t=0.17s (**Fig.4F**, up and down arrows) indicating accumulation of stochastic trial-by-trial diffusion noise^20,21^.

To estimate varCE for the whole population, we estimated constant ϕ in a similar manner for each unit and the resulting population varCE is shown in **Fig.4H**. Population varCE is then a union of units varCE as shown in **Fig.4G**, blue curve. Strong variation of mean spike rates among units may affect the combined population varCE (**Fig.4I**). However, when several major outliers are taken out, the resulting population varCE shows a similar increase during choice period (**Fig.4G**, red curve). Another approach to prevent this variation among the units from contributing to the combined population variance, is to estimate the total variance using the residuals method^20,21^. Residuals were calculated on a trial-by-trial basis, for each unit individually where the mean spike count in each time bin was subtracted from actual spike count across all trials. The population varCE is calculated as a union of varCE for every unit across all trials calculated on residuals instead of raw spike rates (**Fig.4G**, black curve). In all approaches the population varCE exhibit strong increase between t=0.12s and t=0.18s (**Fig.4G**, up and down arrows) consistent with accumulation of stochastic trial-by-trial diffusion noise^20,21^.

### Analysis of corCE

If trial-by-trial variance of spike rates is defined by a diffusion process, then the correlation between pairs of rates at different times during a trial, the CorCE, should increase in consecutive time bins indicating accumulation of noise during evidence accumulation^20,21^. Covariance of conditional expectation (corCE) was calculated on residuals, with a covariance matrix having non-diagonal terms that correspond to corCE while the matrix diagonal corresponds to

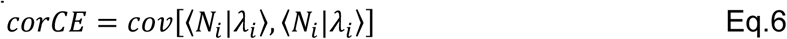

where, 𝑁_𝑖_ is the number of spikes in the bin and 𝜆_𝑖_ the firing rate.

### Streak Index analysis

To analyze local spike count fluctuations we followed the method^22^, that in contrast to the original approach^23^, is calculating the SI for each individual trial and analyzes its distribution across trials that provides a single-trial measure. For that the spike counts from all layers combined in 8 successive 5 ms bins were obtained for each trial. The spike counts in each bin for each trial were compared with the corresponding median spike counts calculated across all trials. The symbol “1” was assigned to bins where the count was higher than the median and “0” was assigned to bins where the count was lower than the median. The resulting binary sequence across trials was used to create a ‘streak index plot’ (**Fig.S10D**). Then we counted how often switching either from 0 to 1 or from 1 to 0 occurred in the resulting 8-digit string to obtain a streak index value for each trial, with the distribution shown in **Fig.4K**. The distribution is close to normal with close to zero mean (0.06±1.36, mean ± s.d., **Fig.4K**) that indicates gradual rather than step-like transition in each trial during ramping.

---

**Figure S1.**
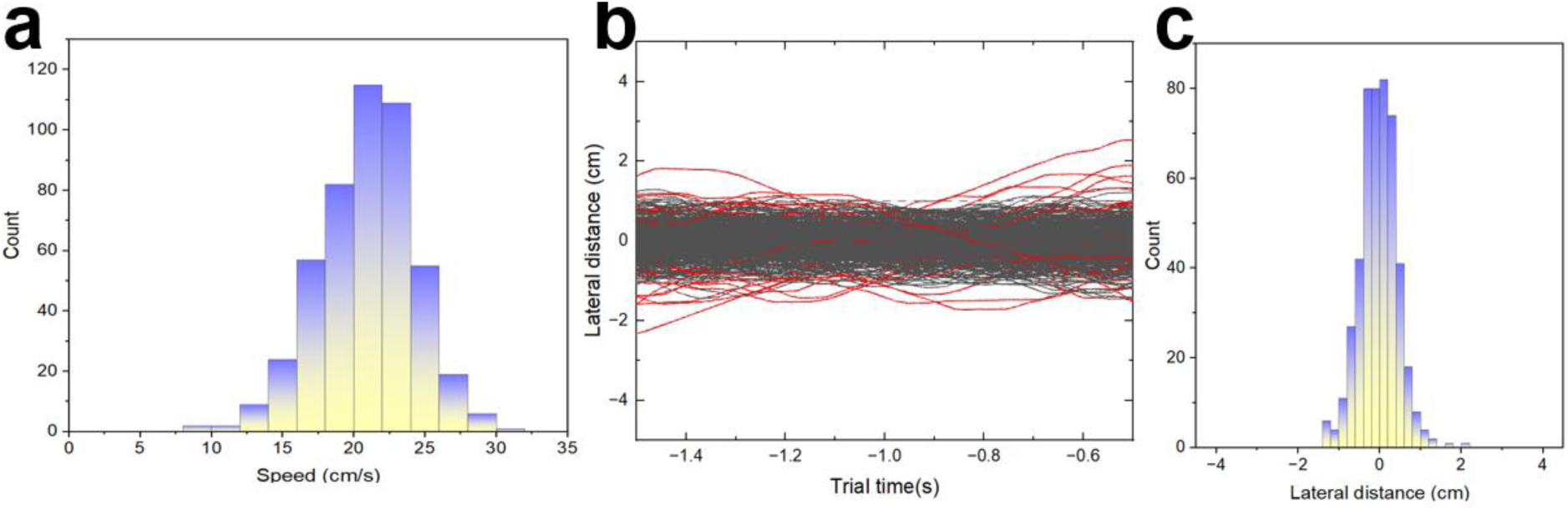
Closed loop straight run. **A)** Histogram of a mouse (animal 503) absolute speed during 1.5-hour 481 trials experiment in Fig.2A. **B)** Mouse trajectories for a closed-loop straight run portion of 481 trials. Red trajectories correspond to deviation of lateral distance larger than 1cm. **C)** Histogram of lateral distance at t = -1s trial time.

**Figure S2.**
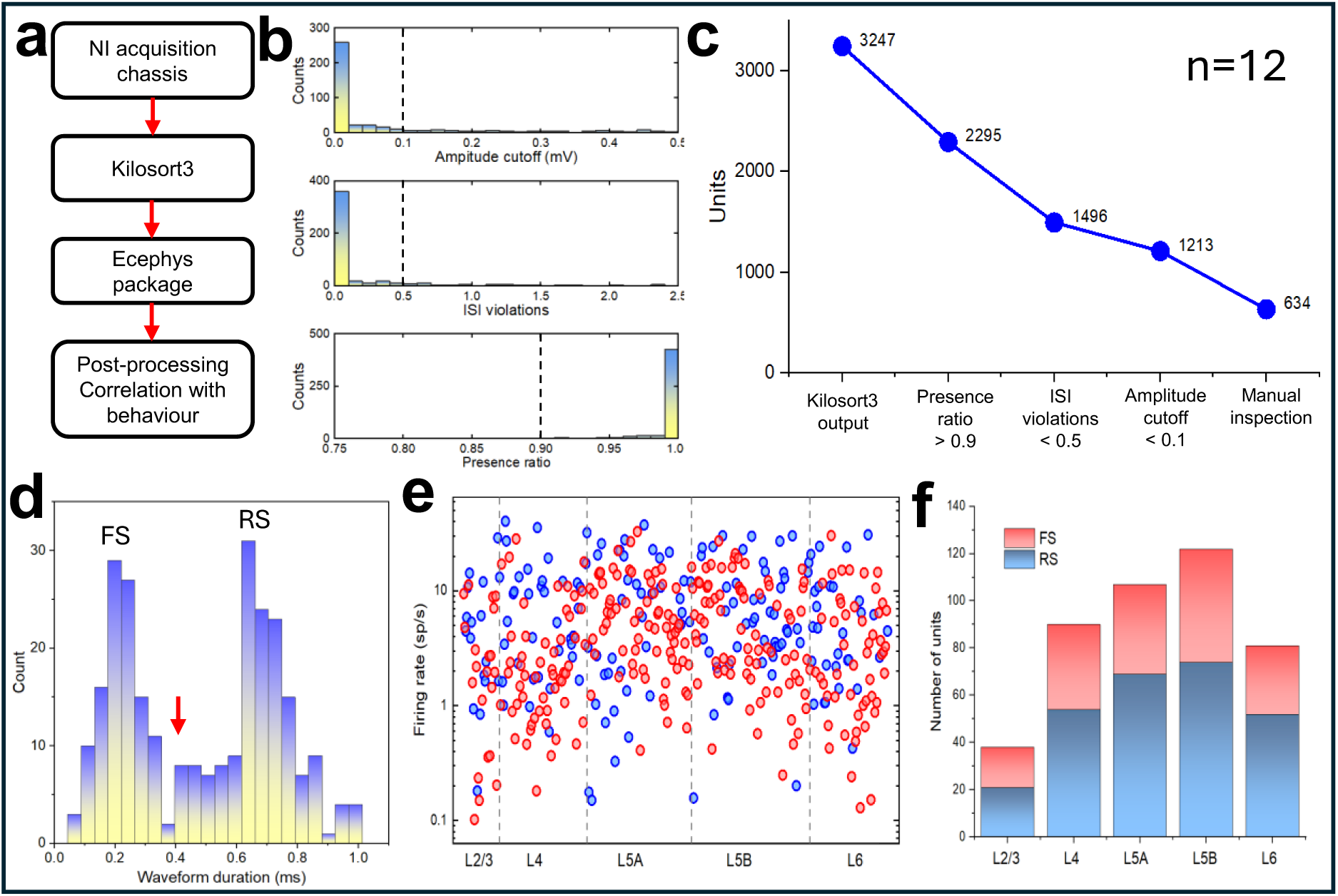
Electrophysiology pipeline. **A)** Block-diagram of spike-sorting and filtering pipeline. **B)** Three major filters to ensure the spike sorting quality: amplitude cutoff (top), ISI violations (middle, and presence ratio (bottom). **C)** Filtered number of units recorded from n=12 animals in the principal whisker C2 barrel (blue) and neighboring barrels (red). See Table S1 for details. **D)** Histogram of sorted spikes waveform duration. Cutoff at 0.4ms was used to classify RS and FS units. **E)** Trial averaged spiking rates for all sorted units ordered by cortical depth and assigned to cortical layers. Red and blue dots correspond to FS and RS units respectively. **F)** Distribution of sorted units across cortical layers for FS (red) and RS (blue) units.

**Figure S3.**
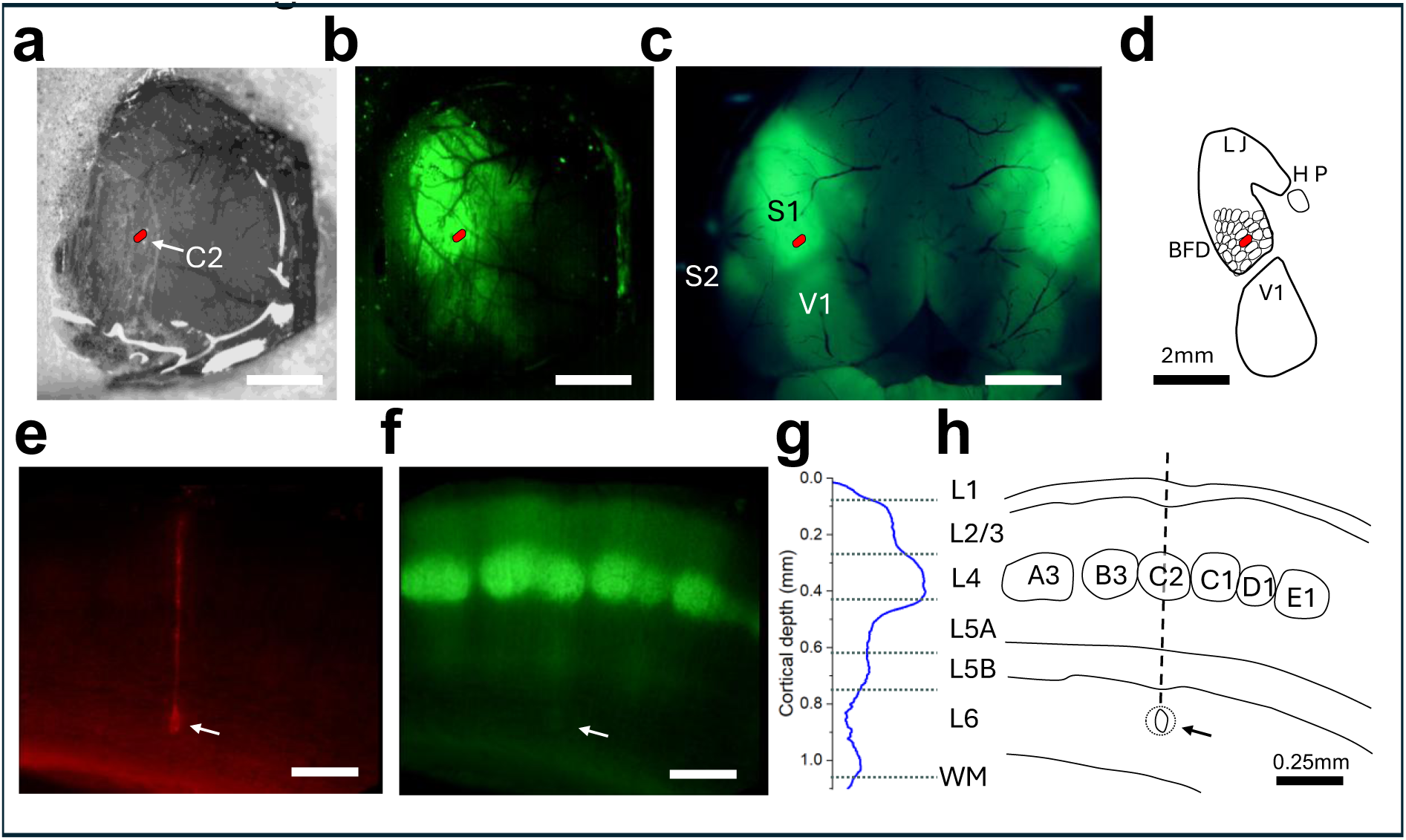
Targeting of C2 barrel and histological verification of probe position. **A)** In-vivo top-down microscope view of the cranial window after surgery at white light illumination. Red oval corresponds to identified position of the principal whisker C2 barrel. **B)** Same as in A) but with fluorescence filter. Green fluorescence from YFP expressed through the barrel cortex is used as a guide for identifying the location of the principal whisker barrel. **C)** Fluorescent microscope image of the brain harvested and partially perfused immediately following the behavior experiment. Partially perfused vasculature pattern is used as a guide to align in-vivo and ex-vivo barrel maps. **D)** Map of the barrel cortex with positions of individual barrels including the principal whisker C2 (red oval) derived from C) and Fig.1G. **E)** Fluorescence micrographs of a coronal slice though a principal whisker barrel column (animal 489) with a green filter showing YFP expressed in layer 4 with projections to layer 2/3 above and layer 5B below. Arrow indicates location of a microlesion produced by the first electrode to identify the location of the probe tip. **F)** Same as in E) but with a red filter showing the probe DiI track. Arrow indicates location of a microlesion performed with the first electrode to identify the location of the probe tip. **G)** Intensity of YFP measured across the column (log scale). **H)** Map of cortical layers and L4 barrels reconstructed from E) and F).

**Figure S4.**
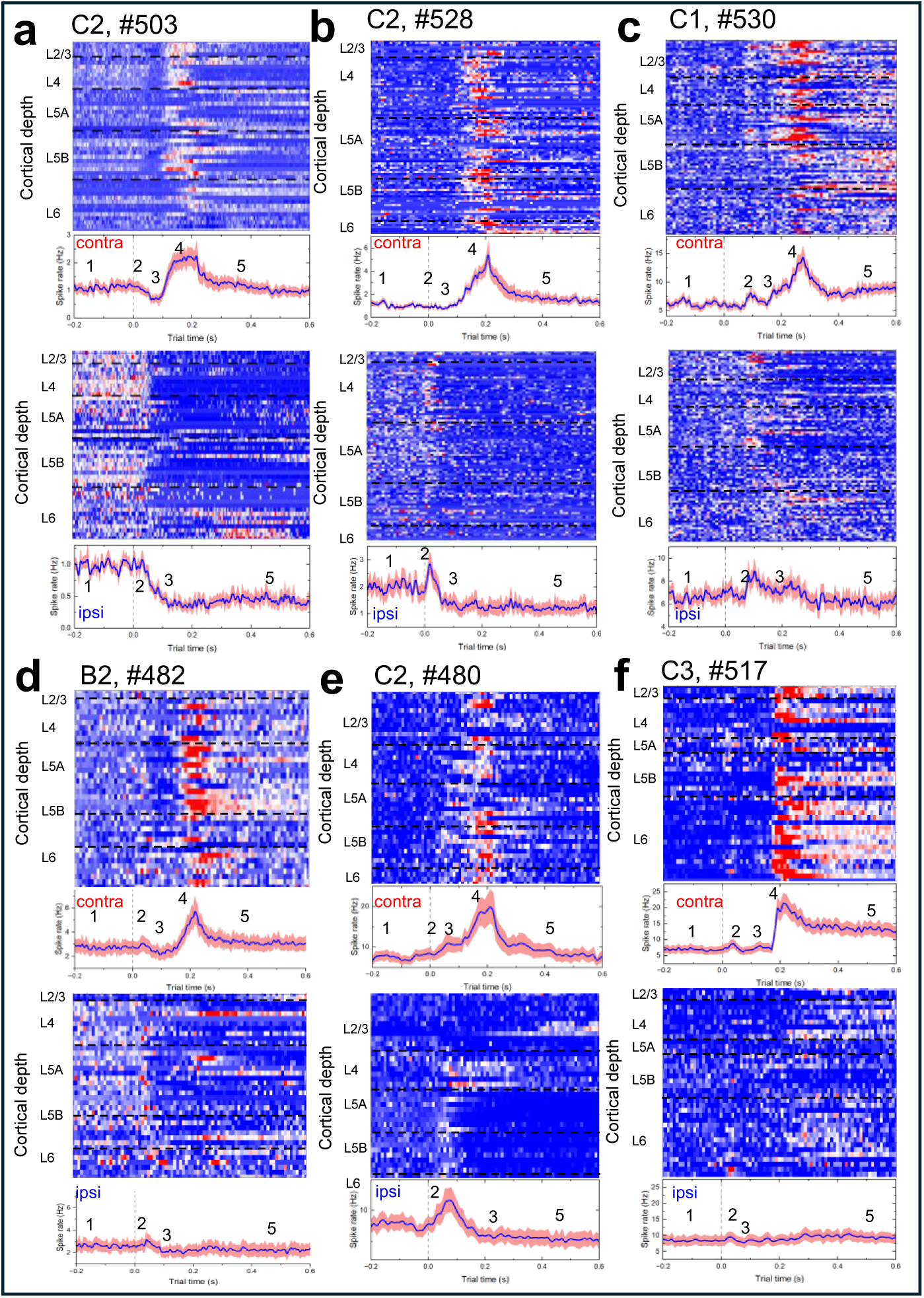
Population neural activity in principal whisker barrel column exhibits strong, staged, and synchronous modulation during decision-making process. **A)** Animal 503. Recording from C2 principal barrel column. **Top panel:** Trial-averaged z-scored spiking activity units across the principal barrel column for OL_L left turns. Units are ordered by cortical depth and assigned to cortical layers. **Second panel**: Population-averaged spike rate for data in A) for OL_L left turns. Numbers correspond to modulation epochs 1 through 5. Blue curve is population mean. Red shadows correspond to mean ± s.e.m. **Third panel**: Same as Top for OL_R right turns. **Bottom panel**: Same as in Second for data in Third panel for OL_R right turns. 48 units, 305 OL_L trials, 304 OL_R. **B)** Animal 528. Recording from C2 principal barrel column. 89 units, 95 OL_L trials, 95 OL_R. **C)** Animal 530. Recording from C1 neighbor barrel column. 113 units, 402 OL_L trials, 403 OL_R. **D)** Animal 482. Recording from B2 neighbor barrel column. 38 units, 350 OL_L trials, 350 OL_R. **E)** Animal 480. Recording from C2 principal barrel column. 42 units, 185 OL_L trials, 190 OL_R. **F)** Animal 517. Recording from C3 neighbor barrel column. 42 units, 270 OL_L trials, 268 OL_R.

**Figure S5.**
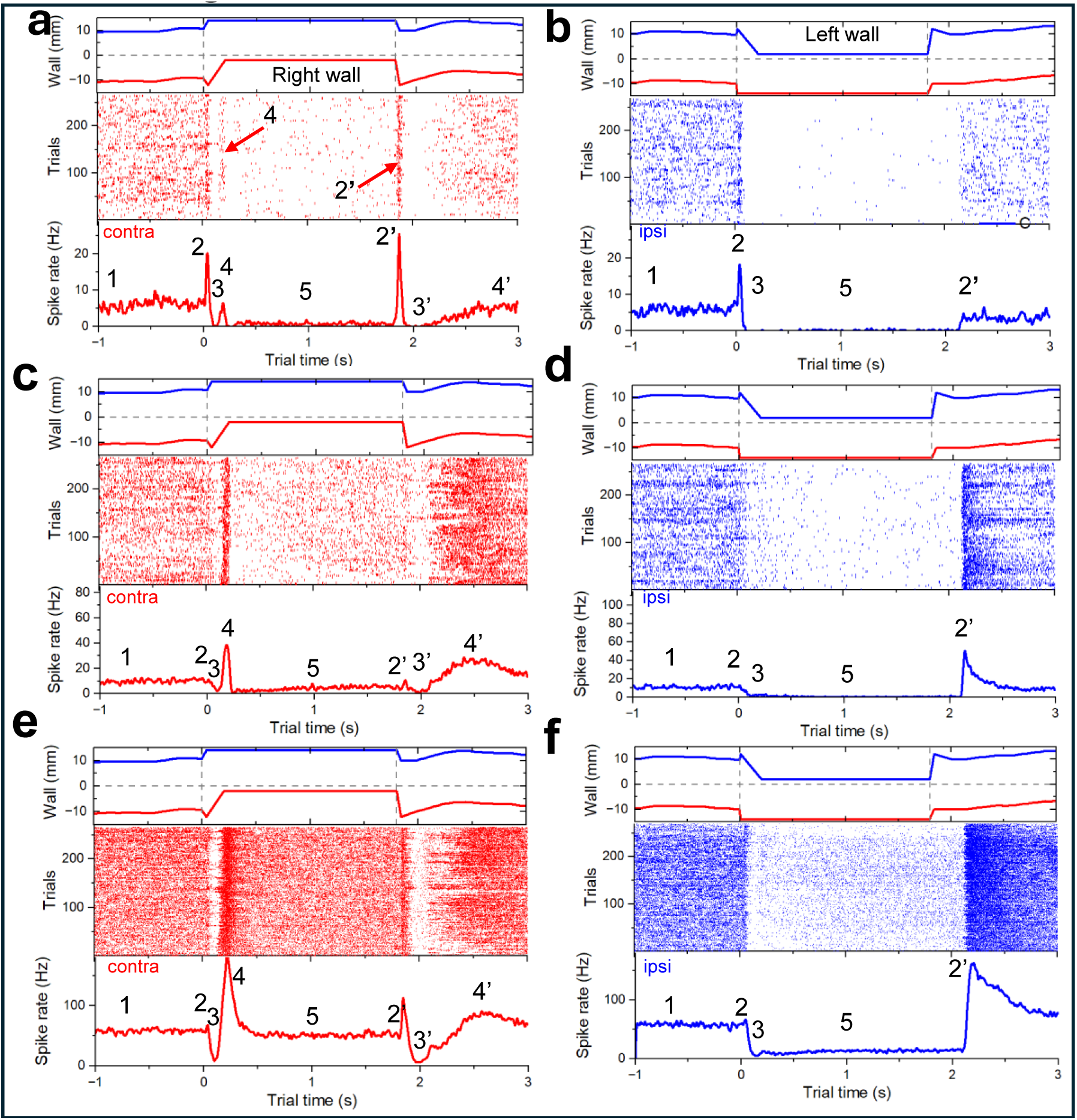
Single unit neural activity in principal whisker barrel column exhibits strong, staged, and synchronous modulation during decision-making process. **A)** Top panel: Right (red) and left (blue) wall positions during the OL_L turn experiment. Middle panel: spiking raster plot of a unit (animal 489, unit 489085, FS, layer 4) recorded during 266 randomized left turns. Bottom panel: Trial-averaged spike rate of a single unit. **B)** Top panel: Right (red) and left (blue) wall positions during the OL_R turn experiment. Middle panel: spiking raster plot of the same unit recorded during 265 randomized right turns. Bottom panel: Trial- averaged spike rate of a single unit. **C)** Top panel: Right (red) and left (blue) wall positions during the OL_L turn experiment. Middle panel: spiking raster plot of a unit (animal 489, unit 489095, RS, layer 5A) recorded during 266 randomized left turns. Bottom panel: Trial-averaged spike rate of a single unit. **D)** Top panel: Right (red) and left (blue) wall positions during the OL_R turn experiment. Middle panel: spiking raster plot of the same unit recorded during 265 randomized right turns. Bottom panel: Trial- averaged spike rate of a single unit. **E)** Top panel: Right (red) and left (blue) wall positions during the OL_L turn experiment. Middle panel: spiking raster plot of a unit (animal 489, unit 489101, RS, layer 5B) recorded during 266 randomized left turns. Bottom panel: Trial-averaged spike rate of a single unit. **F)** Top panel: Right (red) and left (blue) wall positions during the OL_R turn experiment. Middle panel: spiking raster plot of the same unit recorded during 265 randomized right turns. Bottom panel: Trial-averaged spike rate of a single unit. ^6^

**Figure S6.**
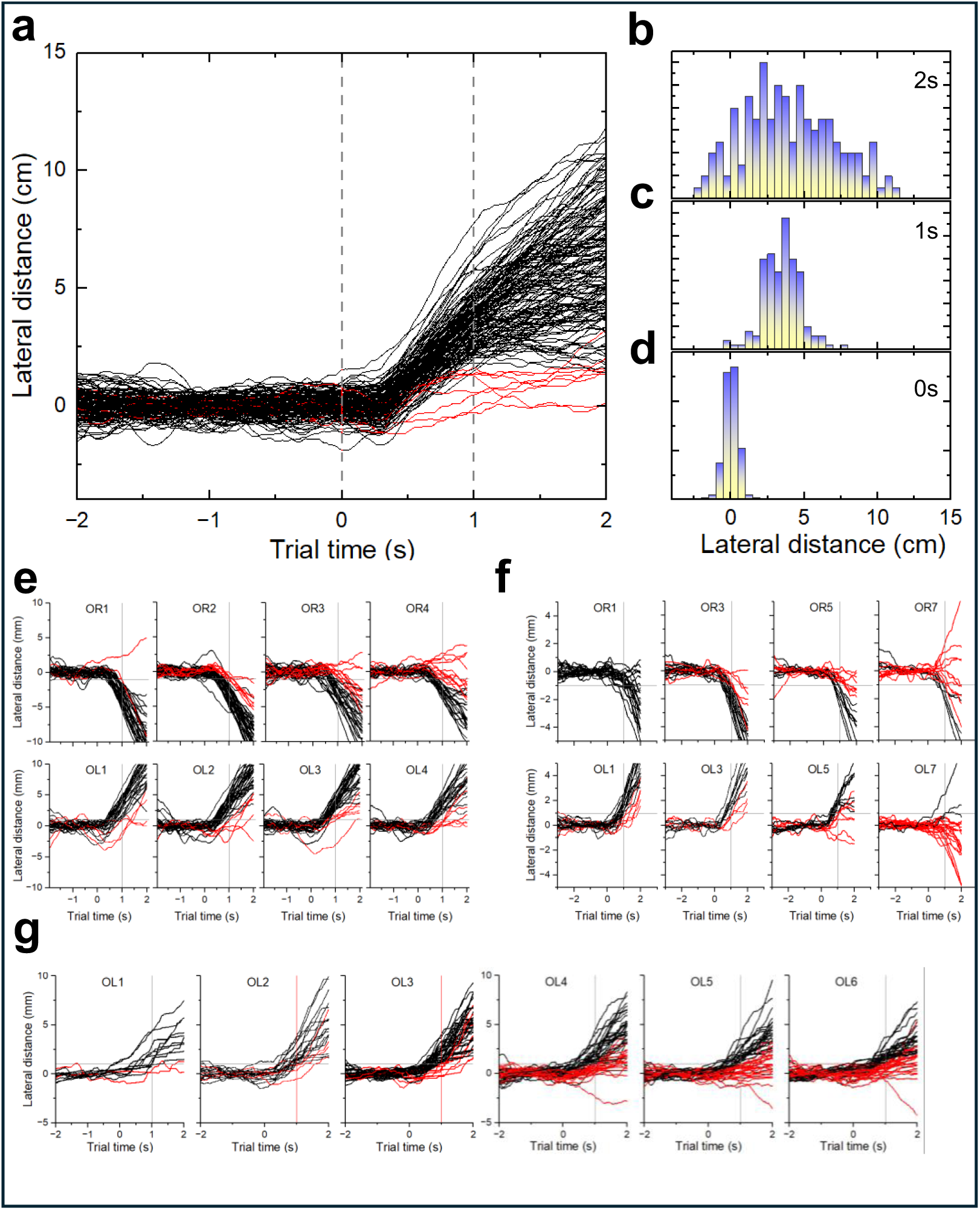
Examples of psychometric performance of different animals. **A)** Trajectories during 520 trials with OL_L left turns (animal 489). Red curves correspond to trials assigned to decision errors. **B, C, D)** Histograms of lateral distance distributions at t=0s (bottom), t=1s (middle), and t=2s (top). While at the beginning of a turn the lateral distribution is narrow, they broaden indicating gradual accumulation of navigation uncertainties. **E)** Top: Trajectories during OL_R right turns (animal 474) for different distances of the wall to the snout. Red curves correspond to trials assigned to decision errors. The closer the wall is approaching the snout the less errors the animal is making. Bottom: Same as on top, but for OL_L left turns. Results are summarized in psychometric curves in Fig.2C, D. **F)** Same as in E) for animal 499. Note very large percentage of errors at the distance of 7mm that correspond to whisker barely touching the wall during foveal whisking. **G)** Same for animal 475 for OL_L left turns.

**Figure S7.**
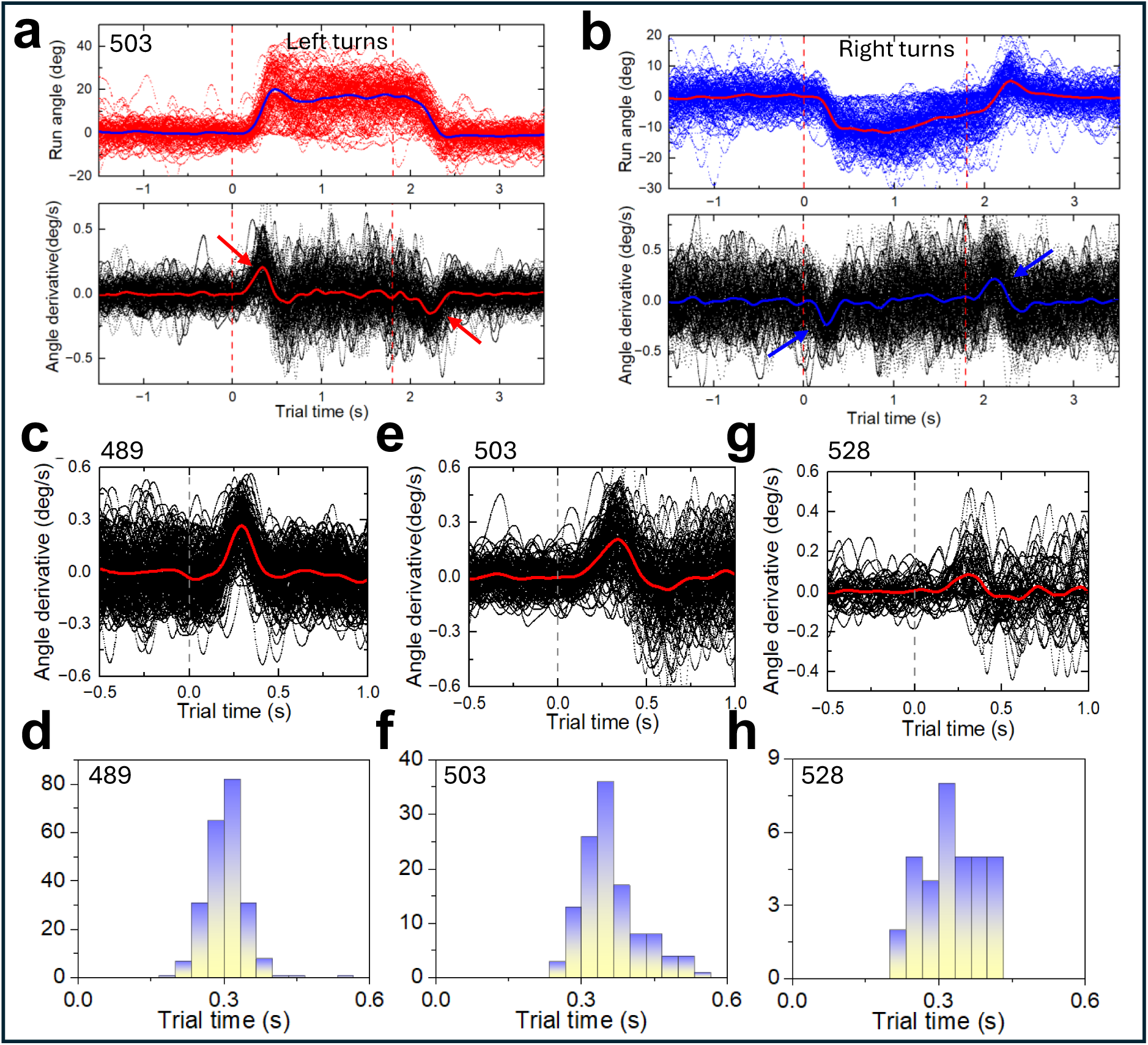
Procedure for extraction of reaction times. **A)** Top: Run angle (animal 503) during a 305 OL_L left turn trials corresponding to data in Fig.2A. Blue curve is ensemble average. **Bottom**: Angle derivative for data in Top. Red curve is ensemble average. Red arrows show characteristic push by RH leg (Fig.2G) at the turn start and LH leg at the turn end. Vertical red dashed lines correspond to the start (t=0s) and the end (t=1.8s) of the turn. **B)** Same as A) but for 304 OL_R right turns randomized with left turns in A). Blue arrows show characteristic push by LH leg (Fig.2G) at the turn start and RH leg at the turn end. Vertical red dashed lines correspond to the start (t=0s) and the end (t=1.8s) of the turn. **C)** Run angle (animal 489) during 520 trials at the start of the left turn corresponding to data in **Fig.S6A**. **D)** Histogram of the maximum angle derivative (reaction time) for data in C). **E)** Run angle (animal 503) during 305 trials at the start of the left turn corresponding to data in D). **F)** Histogram of the maximum angle derivative (reaction time) for data in E). **G)** Run angle (animal 528) during 95 trials at the start of the left turn. **H)** Histogram of the maximum angle derivative (reaction time) for data in G).

**Figure S8.**
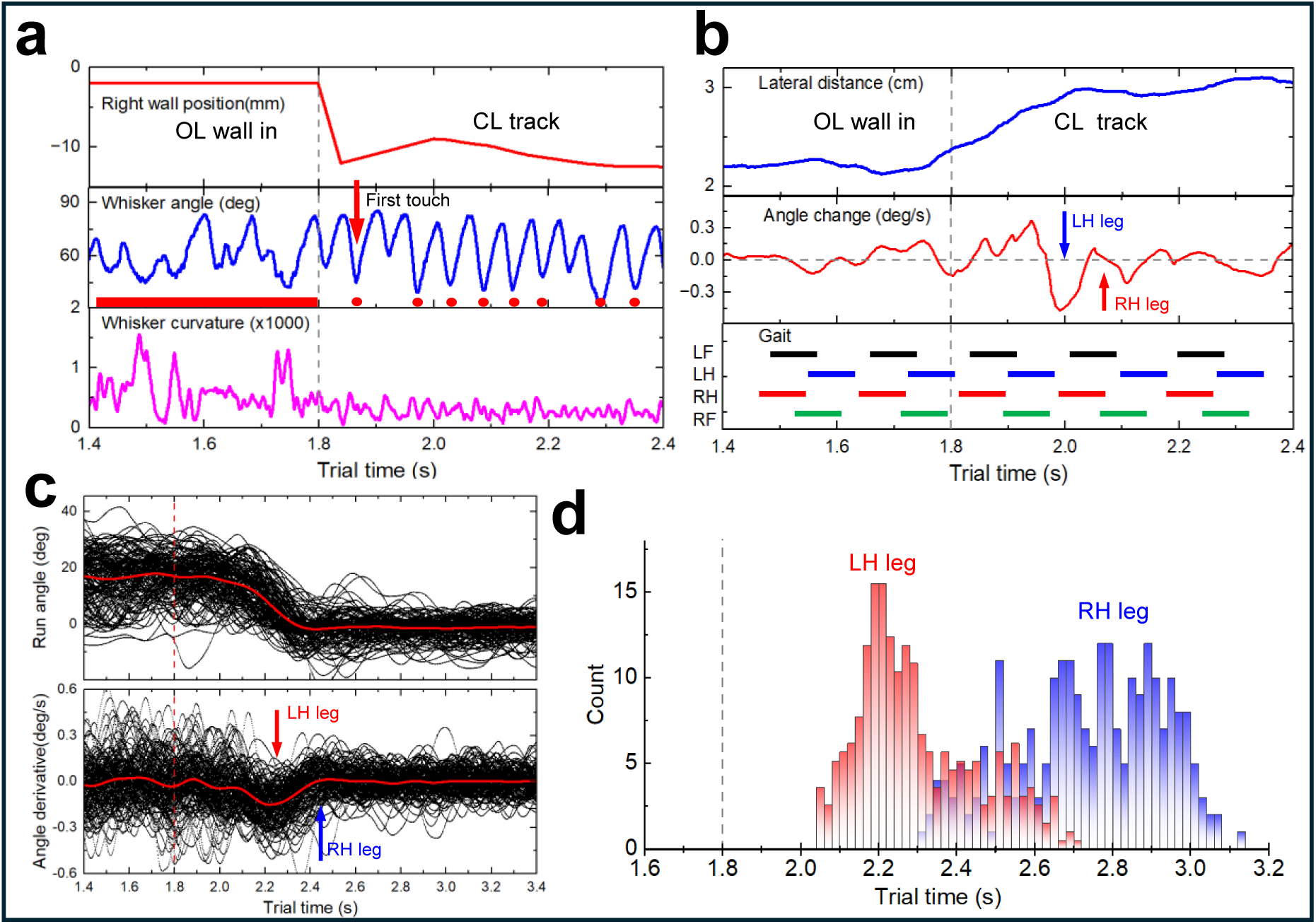
End of the left turn. **A)** Top panel: Position of the right wall at the end of the OL_L left turn (animal 589). Middle panel: extracted whisker angle. Red dots identify moments of whisker touching the wall with the first touch to retracting wall marked by arrow. Bottom panel: Same for extracted whisker curvature. Vertical dotted line corresponds to the start of the wall movement **B)** Top panel: Lateral distance during the left turn trial (animal 589). Middle panel: Angle change curve with a stereotypic sequence of LH leg and RH leg pushes. Bottom panel: tracked gait with left and right hind and fore legs following mixed walking - trotting pattern. **C)** Top panel: Run angle (animal 489) during 520 trials at the end of the left turn. Red curve is ensemble average. Bottom panel: Angle derivative for data in Top. Red curve is ensemble average. Stereotypic sequential pushes by LH leg (red arrow) and RH leg (blue arrow). Vertical dashed line corresponds to the end (t=1.8s) of the turn. **D)** Histogram of the maximum angle derivative (reaction time) for data in C). The decision at the end of the turn is executed not by a single leg push but rather is distributed among several consecutive strides that makes RT analysis complicated. At this extended time, it is also guided by both contra- and ipsi- lateral whiskers as observed with videography and is evident from the appearance of a sharp peak at 2.1s in the ipsi-lateral right turns (Fig.1K).

**Figure S9.**
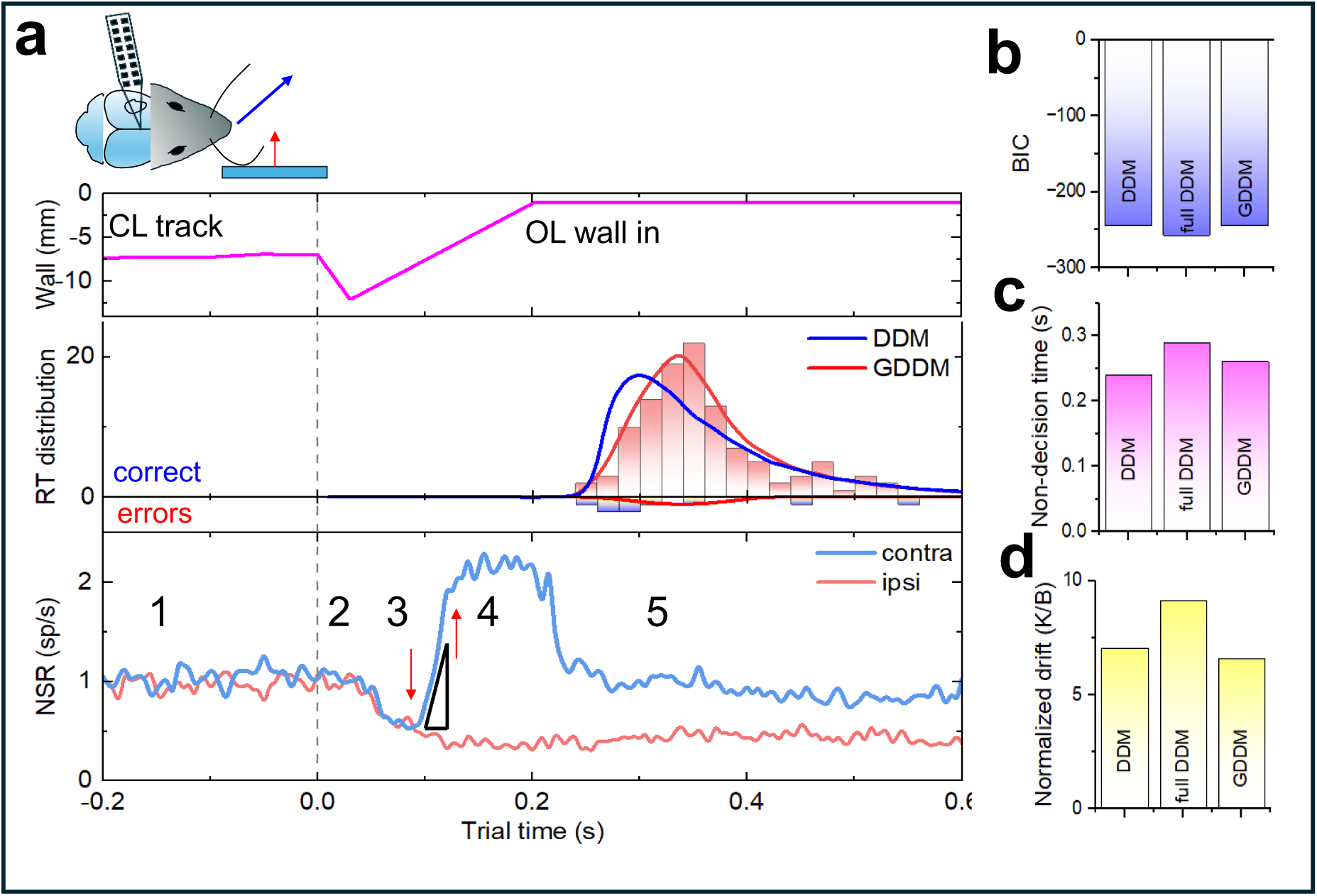
**Ramping neural activity reflects temporal accumulation of sensory evidence.** **A) Top panel**: Position of the right wall during OL_L left turn trials (animal 503). **Middle**: RT distribution fitted by DDM (blue) and Generalized DDM (GDDM, red) models. **Bottom**: Normalized spike rate for contralateral (contra, blue) left and ipsilateral (ipsi, red) right turn trials. Up and down arrows denote times for the beginning and the end of the linear ramping activity. Black triangle shows linear fitting of the ramping slope. **B)** Bayesian information criterion (BIC) for three fitted models – DDM, full DDM and GDDM. **C)** Non-decision time derived from DDM, full DDM, and GDDM fits to the RT. **D)** Drift normalized on bound derived from DDM, full DDM, and GDDM fits to the RT.

**Figure S10.**
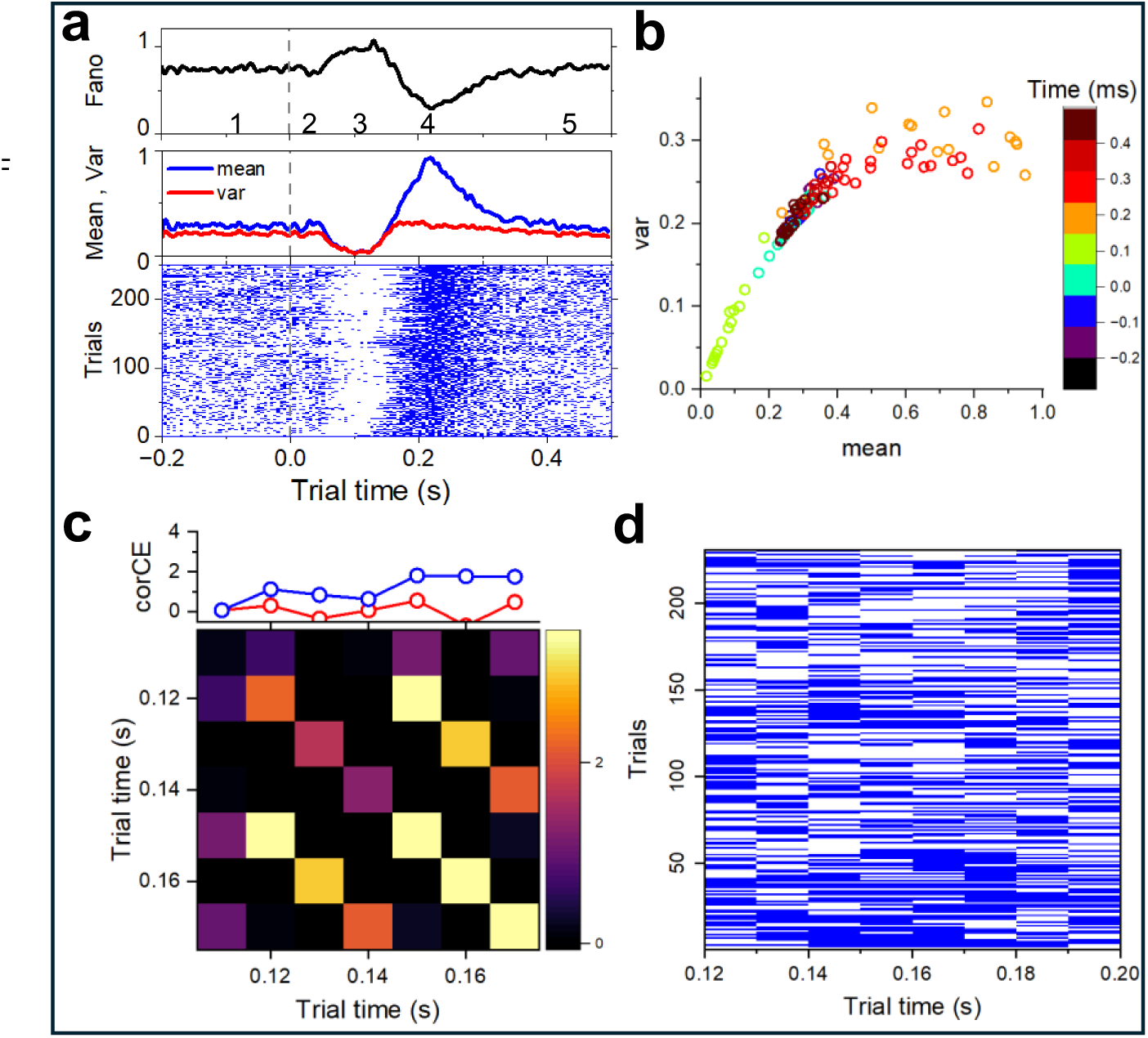
Noise characteristics. **A)** Top panel: Fano factor for a single unit (489101, RS, layer 5A, animal 489) recorded during OL_L left turns. Middle panel: Mean (blue) and variance (red) for a single unit (489101, RS, layer 5A, animal 489). Bottom panel: Raster plot for a single unit (489101, RS, layer 5A, animal 489). **B)** Dependence of spiking variance (var) on mean spiking rate (mean) for the same unit. **C)** Covariance matrix CorCE calculated on residuals when bins are randomly shuffled. Top: corCE values along the matirx diagonal (red) and along the top row (blue) when bins are randomly shuffled. **D)** Streak matrix for data in Fig.5K.

**Figure S11.**
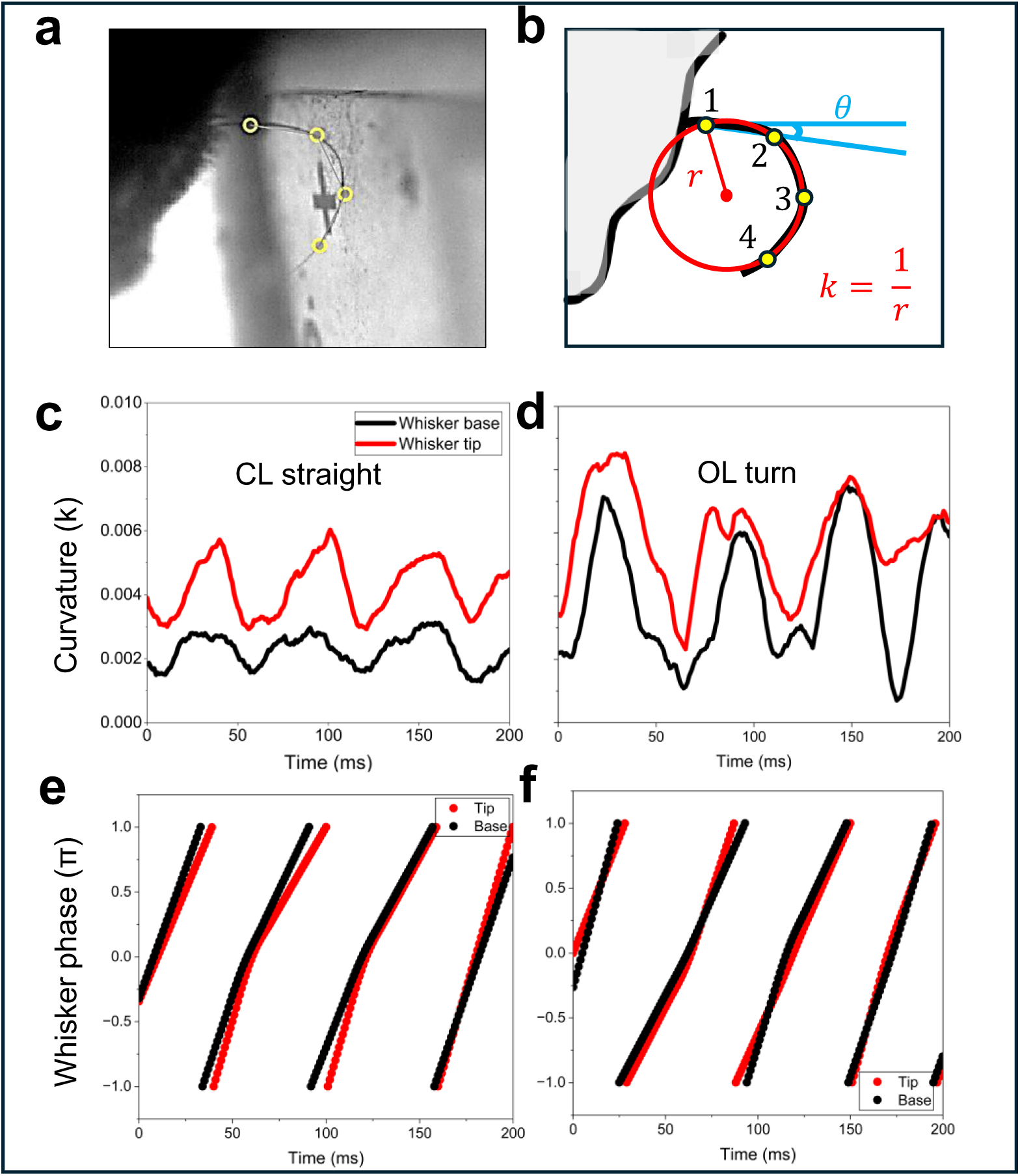
Whisker curvature calculations at base and tip. **A)** Example frame from 1000fps whisker tracking camera, showing four tracked points along the whisker as yellow circles from SLEAP based pose estimation model (Methods). **B)** Schematics detailing calculation of whisker parameters. Circle is fit to either three points closest to the whisker base, or the whisker tip, with curvature (k) calculated as the inverse of the radius. Whisker angle (θ) is calculated from the perpendicular line from the mouse’s face using the two points closest to the whisker base. **C)** Comparison of whisker curvature calculated at the whisker base, or the whisker tip during example 200ms segment CL straight run. **D)** Comparison of whisker curvature calculated at the whisker base, or the whisker tip during example 200ms segment OL turn. **E)** Phase of whisking cycle for the same 200ms segment of CL straight run as in C) calculated for both whisker base and tip. **F)** Phase of whisking cycle for the same 200ms segment of OL turn as in D) calculated for both whisker base and tip.

**Figure S12.**
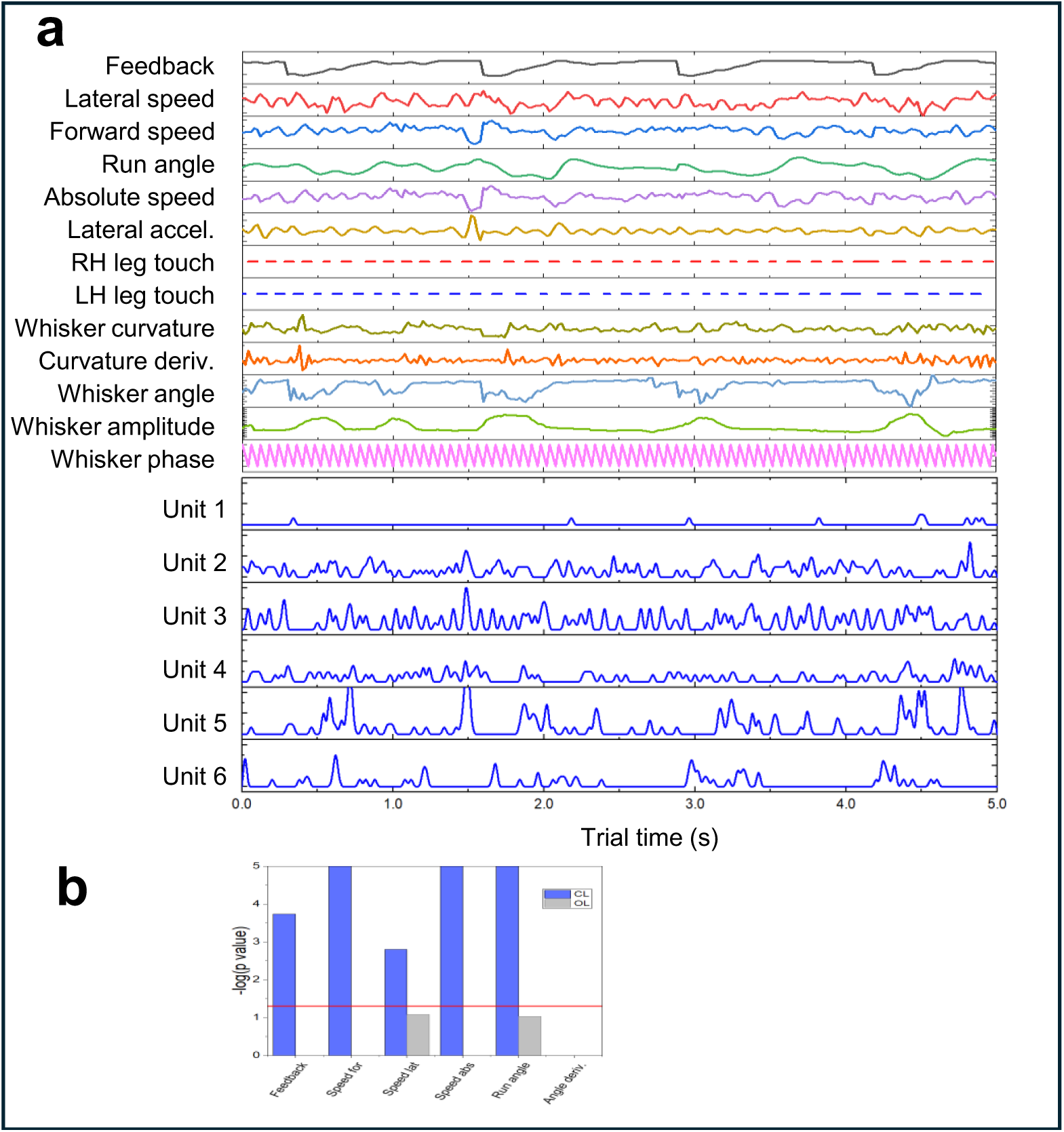
Variables encoded during decision. **A)** Example 5s segment of CL straight running showing precise alignment of 13 behavioral variables used for GLM design matrix construction with 5 example neurons organized into 20ms time bins. **B)** Example unit where separate GLMs were constructed using CL straight periods and OL turn decisions. 100 trials for each condition with 2s segment extracted from each trial. Log likelihood ratio (Methods) shows unit encoding variables preceding but not during decision.

**Video S1.**
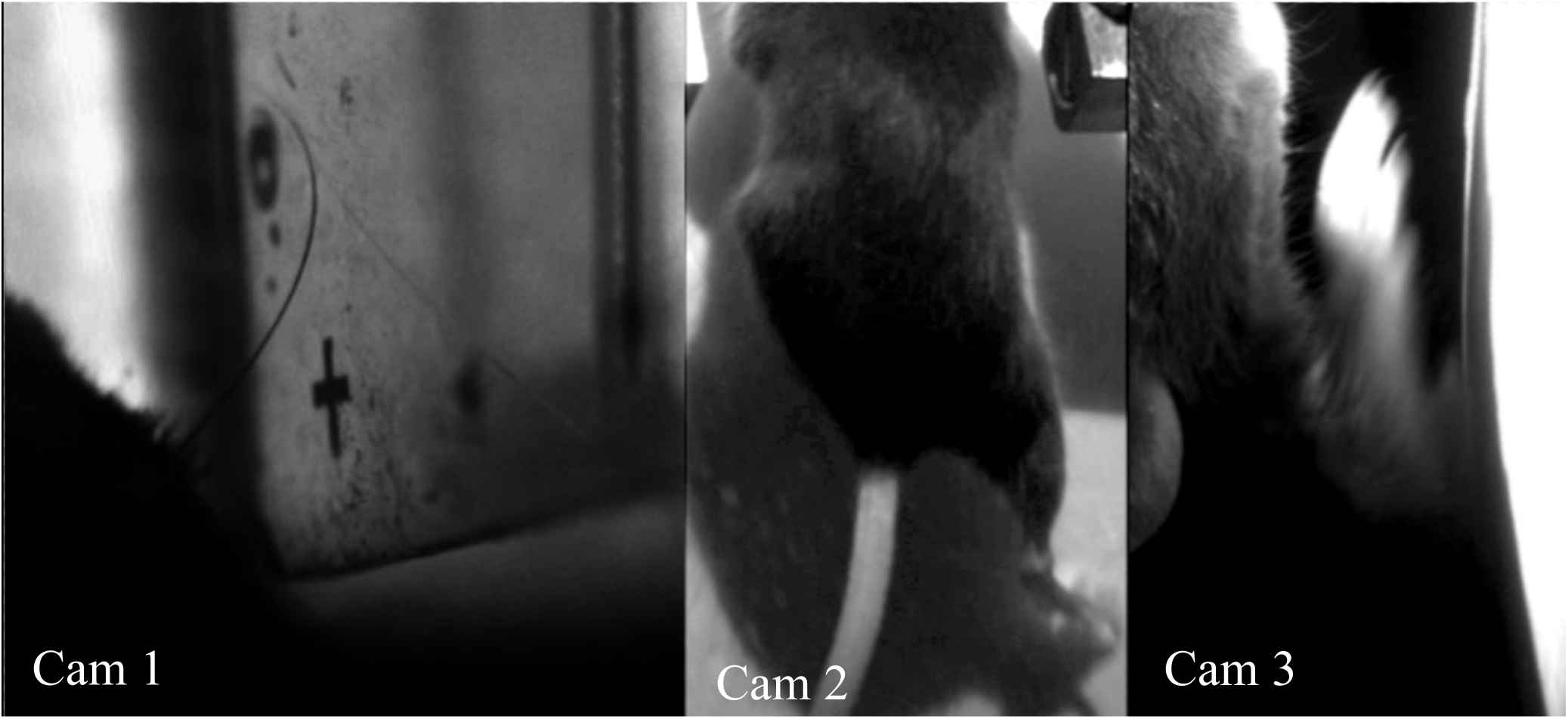
Synchronized recordings of whisker, fore and hind legs. The video combined with 3 cameras showing synchronized video streams from whisker recording (cam 1) together with recordings of hind (cam 2) and fore (cam 3) legs during whisker-guided navigation.

**Table S1.**
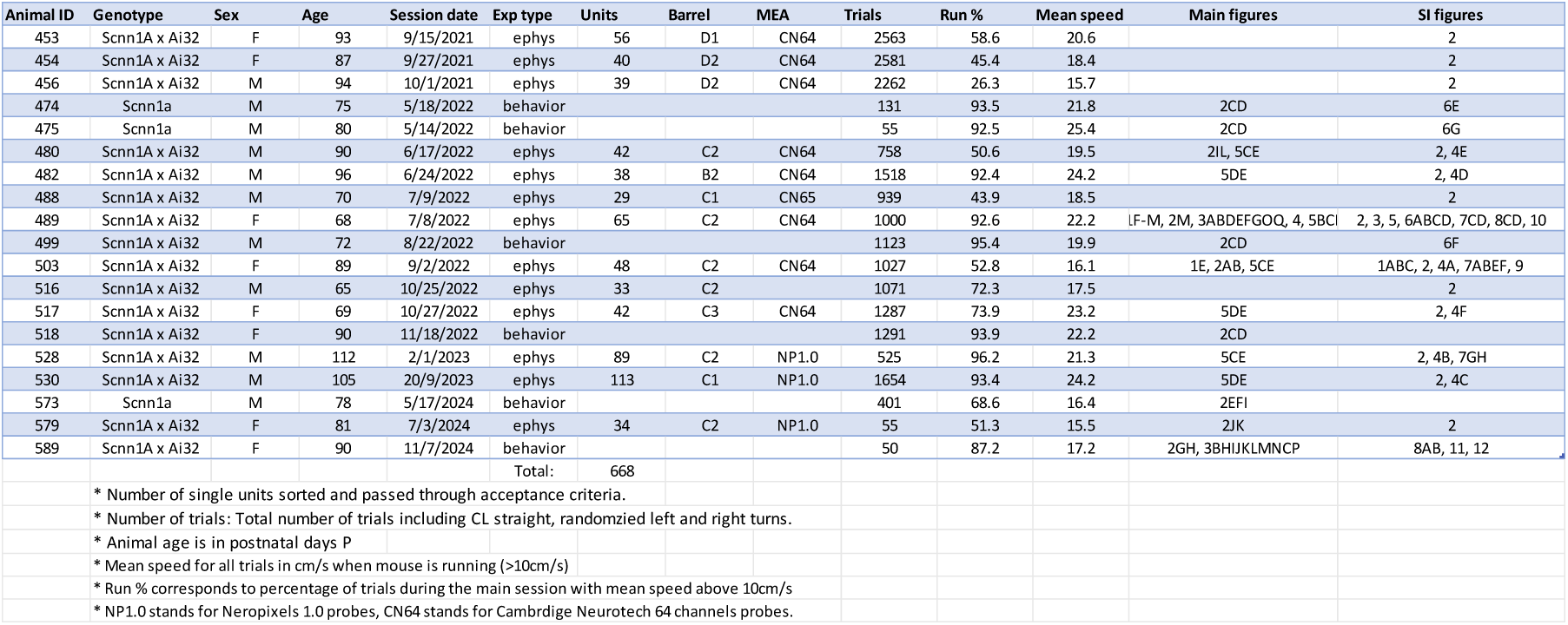
List of mice used for behavioral and electrophysiology experiments.

**Table S2.** Parameters of models best fitted to RT data Fig.3B (animal 489)

| Parameter | DDM | full DDM | GDDM |
| --- | --- | --- | --- |
| Animal | 489 | 489 | 489 |
| drift | 2.91 | 2.81 | 4.04 |
| noise | 0.25 | 0.1 | 0.62 |
| B | 0.59 | 0.54 | 0.6 |
| x0 |  | 0.01 | 0.14 |
| xo width |  | 0.04 | 0.01 |
| tind | 0.1 | 0.11 | 0.15 |
| tind width |  | 0.05 | 0.09 |
| leak |  |  | 93.68 |
| tau |  |  | 10.53 |
| drift/B | 4.93 | 5.20 | 6.73 |
| BIC | -664 | -666 | -690 |

**Table S3.** Parameters of models best fitted to RT data Fig.S9 (animal 503)

| Parameter | DDM | full DDM | GDDM |
| --- | --- | --- | --- |
| Animal | 503 | 503 | 503 |
| drift | 2.54 | 1.46 | 2.77 |
| noise | 0.75 | 0.48 | 0.87 |
| B | 0.36 | 0.16 | 0.42 |
| x0 | 0.01 | 0.04 | 0.03 |
| xo width |  | 0.01 | 0.04 |
| tind | 0.24 | 0.29 | 0.26 |
| tind width |  | 0.09 | 0.26 |
| leak |  |  | 1.15 |
| tau |  |  | 2.31 |
| drift/B | 7.06 | 9.13 | 6.61 |
| BIC | -244 | -257 | -244 |

