## Supplementary figures and images for "Neural Correlates of Perceptual Decision Making in Primary Somatosensory Cortex"

### Video S1

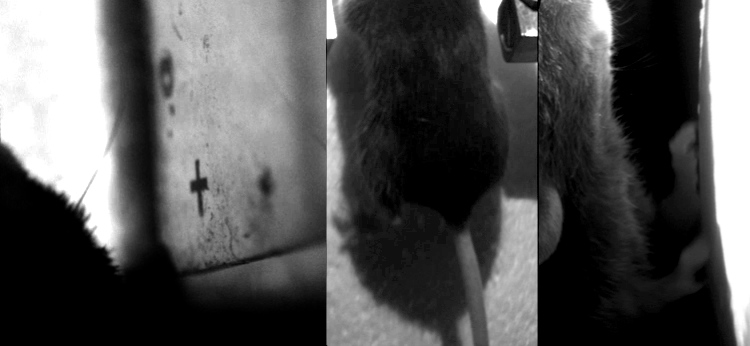
